# Human cortical mosaicoids reveal impact on spatial organization and progressive loss of cross-genotype functional coupling in MECP2 mosaicism

**DOI:** 10.64898/2026.11.11.765820

**Authors:** Tatsuya Osaki, Gregg R. Heller, Mriganka Sur

**Affiliations:** Picower Institute for learning and memory, Massachusetts Institute of Technology, Cambridge, MA 02139, USA; Whitehead Institute for Biomedical Research, Massachusetts Institute of Technology, Cambridge, MA 02139, USA; Department of Intelligent Systems Engineering, Indiana University, Bloomington, IN, 47408, USA; Department of Brain and Cognitive Sciences, Massachusetts Institute of Technology, Cambridge, MA 02139, USA

## Abstract

Cellular mosaicism arising from postzygotic mutation or X-chromosome inactivation is a major determinant of phenotypic variability in neurodevelopmental disorders. Rett syndrome, which is caused by mutations in the X-linked gene MECP2, is a striking example in which wild-type and mutant neural cells coexist within the developing brain. However, how MECP2 mosaicism shapes human cortical tissue organization and circuit assembly remains poorly understood. Here, we developed MosaicoidX, a human mosaic organoid platform combining controlled co-seeding of isogenic wild-type and MECP2-mutant hiPS cells with genotype-resolved calcium imaging, computational modeling, and single-cell transcriptomics. FACS-based single-cell seeding enabled reproducible generation of cortical mosaicoids with defined initial genotype ratios. Structural two-photon imaging revealed mutation-dependent spatial organization, with MECP2[V247X] mosaicoids (harboring a truncating mutation) showing pronounced skewing of radial and angular segregation compared with MECP2[R306C] mosaicoids (harboring a missense mutation). MosaicoidSim, a computational model, showed that genotype-dependent adhesion, proliferation, differentiation, and migration parameters can account for the emergence of mosaic spatial patterning. Genotype-resolved two-photon calcium imaging demonstrated asymmetric functional connectivity, in which wild-type neuron pairs formed stronger and more frequent activity-coupled interactions than mutant-mutant or wild-type-mutant pairs. Finally, single-cell RNA-seq combined with ligand-receptor analysis inferred genotype- and cell-type-dependent alterations in synaptic signaling pathways, including GABAergic and neurexin-neuroligin interactions. Together, these results show that MECP2 mosaicism actively shapes organoid development through both cell-autonomous and non-cell-autonomous mechanisms. MosaicoidX provides a general framework for dissecting genetic mosaicism in Rett syndrome and other mosaic neurodevelopmental disorders.

## Introduction

Cellular mosaicism, whether caused by postzygotic *de novo* mutations or random X-chromosome inactivation (XCI), is a central factor shaping phenotypic variability and disease severity in neurodevelopmental disorders. Rett syndrome (RTT) is a rare neurodevelopmental disorder that is primarily caused by *de novo* germline mutations of methyl-CpG binding protein 2 (MeCP2) on X-chromosome, and predominantly affects girls^1–3^. Disease severity can vary significantly between individuals due to the many variations of MeCP2 mutations^4–7^ and genomic mosaicism by random X-chromosome inactivation (XCI)^8,9^, representing an additional layer of variable symptoms and effects^10–12^. Heterogeneity by mosaicism and regional MeCP2 expression ^13–16^ is one of the critical reasons for diversity in the severity of RTT^17^, although XCI ratio itself, confirmed by blood, does not impact phenotypic severity^18^. While the location of MeCP2 mutations influences differential molecular mechanisms^19^, it alone does not fully explain the clinical effects of RTT as the severity of the disease varies widely among individuals with the same MeCP2 mutation^17,20–22^. In addition, disease severity may depend not just on the global WT/Mut ratio, but on how these populations interconnect with each other and are spatially organized in the local circuit. Because cortical function depends on columnar organization ^23–27^, mutant cells clustered within specific columns should cause more severe dysfunction than a balanced distribution. This spatial patterning and co-development of mutant and wild-type cellular states in the brain, together with asymmetric synapse formation may critically shape neural development, yet its impact on mosaic circuit formation remains poorly understood and its mechanism has never been directly tested in developing human tissue^28^. Furthermore, while some variability could be due to skewed XCI^13–16^, individuals with balanced XCI have substantially different disease phenotypes^17,22^, suggesting that other genetic or environmental factors strongly influence the clinical severity of RTT, where individuals with a higher proportion of cells expressing the mutated *MECP2* gene typically experience a more severe phenotype^16^. Despite the critical importance of mosaicism, RTT mouse models using heterozygous females are limited in their ability to resolve cell-type-specific contributions, especially the communication between wild-type and mutant neurons during development.

Cortical organoids from human induced pluripotent stem cells (hiPS cells) have emerged as powerful experimental systems for their ability to recapitulate early human neurodevelopmental processes^29,30^. This *in vitro* approach allows us to expand our current understanding of cortical development, which is mostly constructed from studies with human individuals, animal experiments with rodents, and conventional 2D cell cultures *in vitro*. Importantly, patient-derived organoids facilitate analyses of pathological mechanisms underlying not only anatomical alterations^31–33^, but also their neuronal functionality and cortical developmental dynamics^19,32–35^. We have previously demonstrated differential impact of two different MeCP2 mutations with RTT patient-derived cortical organoids, where we found altered cortical development and functional connectivity with altered network structures, characterized by changes in the local clustering coefficient and small-world propensity (SWP) ^19,36^. However, these organoid models do not consider heterozygous conditions similar to that seen in RTT human patients. Because RTT primarily affects females, random XCI results in a cellular mosaic state^8,10–18,20–22^, where the brain contains a mix of healthy and mutant cells. The variable ratio of these two cell types is thought to directly influence disease severity, but this has never been directly examined in an organoid model.

Although simple mixing of genetically distinct iPS cells into organoids and chimeroids^37–40^ from different healthy donors has been demonstrated to normalize interindividual genetic variation, critical questions remain: (1) what methodological principles govern successful mosaic organoid generation, and (2) how does mosaicism functionally impact early developmental processes, particularly when modeling *de novo* mutations in Rett syndrome. Previous studies^41^ using somatic mosaic midbrain organoids composed of DEPDC5^+/−^ and DEPDC5^−/−^ cells have provided important insights into epilepsy, demonstrating that even a relatively small proportion of mutant neurons are sufficient to induce epileptiform activity in a mosaic context. However, further analyses are needed to define genotype- and cell-type-specific molecular pathways and to understand how mosaic context influences neuronal network assembly.

Here, we establish MosaicoidX, a platform combining controlled co-differentiation of mutant and isogenic iPS cell-derived cortical organoids with longitudinal multi-modal analysis to test whether MECP2 mutations disrupt neuronal network balance through both cell-autonomous and non-cell-autonomous mechanisms when present in mosaic compositions. FACS-based controlled single-cell seeding enabled reproducible generation of homogeneous mosaicoids with defined initial genotype composition. In addition, by combining Mosaicoid two-photon imaging with CRISPR-based fluorescent labeling, we preserve genotype information while quantifying circuit-level organization at single-cell resolution. This integrated approach demonstrates that mosaic MECP2 mutations disrupt network assembly through genotype-dependent cell-autonomous dysfunction and the emergence of asymmetric functional connectivity patterns between mutant and wildtype populations. In parallel, we developed MosaicoidSim, a computational framework that simulates organoid growth and cellular migration in mosaic contexts. By systematically varying initial seeding ratios, genotype-specific cell adhesion properties, and proliferation/differentiation dynamics, MosaicoidSim predicts how these parameters collectively determine final cell distribution, spatial organization, and mosaic composition. Furthermore, MosaicoidSeq, integrating single-cell RNA-seq with genotype demultiplexing followed by NeuronChat analysis, identifies genotype- and cell-type-dependent alterations in synaptic development and intercellular signaling that underlie asymmetric connectivity patterns in mosaicoids. These multi-modal analyses demonstrate that cellular mosaicism is not a passive admixture but an active developmental determinant that sculpts tissue organization, circuit assembly, and emergent cellular phenotypes through integrated cell-autonomous and non-cell-autonomous mechanisms. More broadly, MosaicoidX establishes cortical mosaicoids as a versatile and experimentally tractable platform in the organoids field for dissecting the neurodevelopmental consequences of XCI in Rett syndrome and other X-linked disorders. By enabling prospective control of genotype ratios, longitudinal tracking of cellular behaviors, and multi-scale analysis spanning imaging, computation, and single-cell transcriptomics, this platform provides a general strategy to investigate how mosaicism contributes to phenotypic heterogeneity, variable disease severity, and selective cellular vulnerability in the developing human brain.

## Results

### Controlled single-cell seeding generates reproducible human cortical mosaicoids with defined genotype composition

Mosaicism is a fundamental characteristic of Rett syndrome and can influence the severity of symptoms through factors such as tissue specificity, XCI in post-zygote, epigenetics, and environmental influences (**Fig. 1A**). While the majority of Rett syndrome patients exhibit balanced XCI^18^, prior studies have identified instances of *de novo* mosaic C>T nonsense mutations characterized by approximately 25% abnormal cells^14,42^, resulting in patients displaying the full spectrum of Rett syndrome phenotypes despite undominant mosaic mutations. To recapitulate this mosaicism *in vitro* with the missense R306C mutation and truncated V247X mutation, we established Rett syndrome mosaic organoids ^43^(Mosaicoids) from human female iPS cells with MeCP2 mutations on Xa and isogenic control pair. First, mutant iPS cells were labeled with EGFP and isogenic control with tdTomato by CRISPR/Cas9 knock-in into the AAVS1 safe harbor locus (**Fig. 1B**). Targeted insertion of fluorescent transgenes into the AAVS1 locus ensures stable, uniform expression without disrupting endogenous genes. To generate RTT cortical mosaicoids, we employed fluorescence-activated cell sorting (FACS) to precisely control the seeding of individual iPS cells from wildtype and MECP2-mutant lines into 96-well plates at defined ratios, providing three critical advantages: (1) elimination of cell doublets and multiples for true single-cell seeding; (2) precise quantitative control of cell number and genotypic composition; and (3) verification of fluorescent marker expression prior to organoid initiation. Sorted single-cell suspensions were cultured through neural induction and subsequently differentiated into cortical organoids. (**Fig. 1C**). Controlled FACS-based seeding combined with ROCK inhibitor (ROCKi) treatment generated spatially homogeneous mosaicoids (**Fig. 1D**(i)), whereas uncontrolled seeding produced distinct phenotypes: ROCKi-treated cultures exhibited high spatial heterogeneity with variable clustering patterns (**Fig. 1D (ii)**), while untreated cultures underwent genotype-dependent segmentation (**Fig. 1D (iii))**. These findings support the hypothesis that true-single-cell seeding and initial spatial geometry are fundamental determinants of organoid development (**Fig. 1F, G**) and genotype-dependent differences in adhesion may contribute to segmented mosaic organization. Notably, the mosaicoids using human iPS cells which have different genotypes (donor) showed enhanced segmentation, indicating that genetic variation alters adhesive properties and promotes spatial separation (**Fig. S1A**). Mosaicoid growth and cell composition was monitored longitudinally by characterizing the population change of WT/mutant cells in mosaicoids by flow cytometry (**Fig. 1G**). Interestingly, MeCP2[V247X] mosaicoids showed skewed growth of mutant cells which reached to 70% of the population while MeCP2[R306C] mosaicoids showed 50%/50% uniform growth even after 3 months of culture (**Fig. 1F**), implying in typical MeCP2 mutation represented balanced ratio. This finding is explained by cell-autonomous effects via the differential growth rate between wildtype and mutant cells because this can be also observed in the 100% control organoid and 100% mutant organoids (**Fig. S1B**). Controlled cell seeding with ROCKi allowed homogenous mosaicoids after 3 months (**Fig. S1D**), while seeding without ROCKi resulted in heterogenous mosaicoids (**Fig. S1C**).

**Figure 1.**
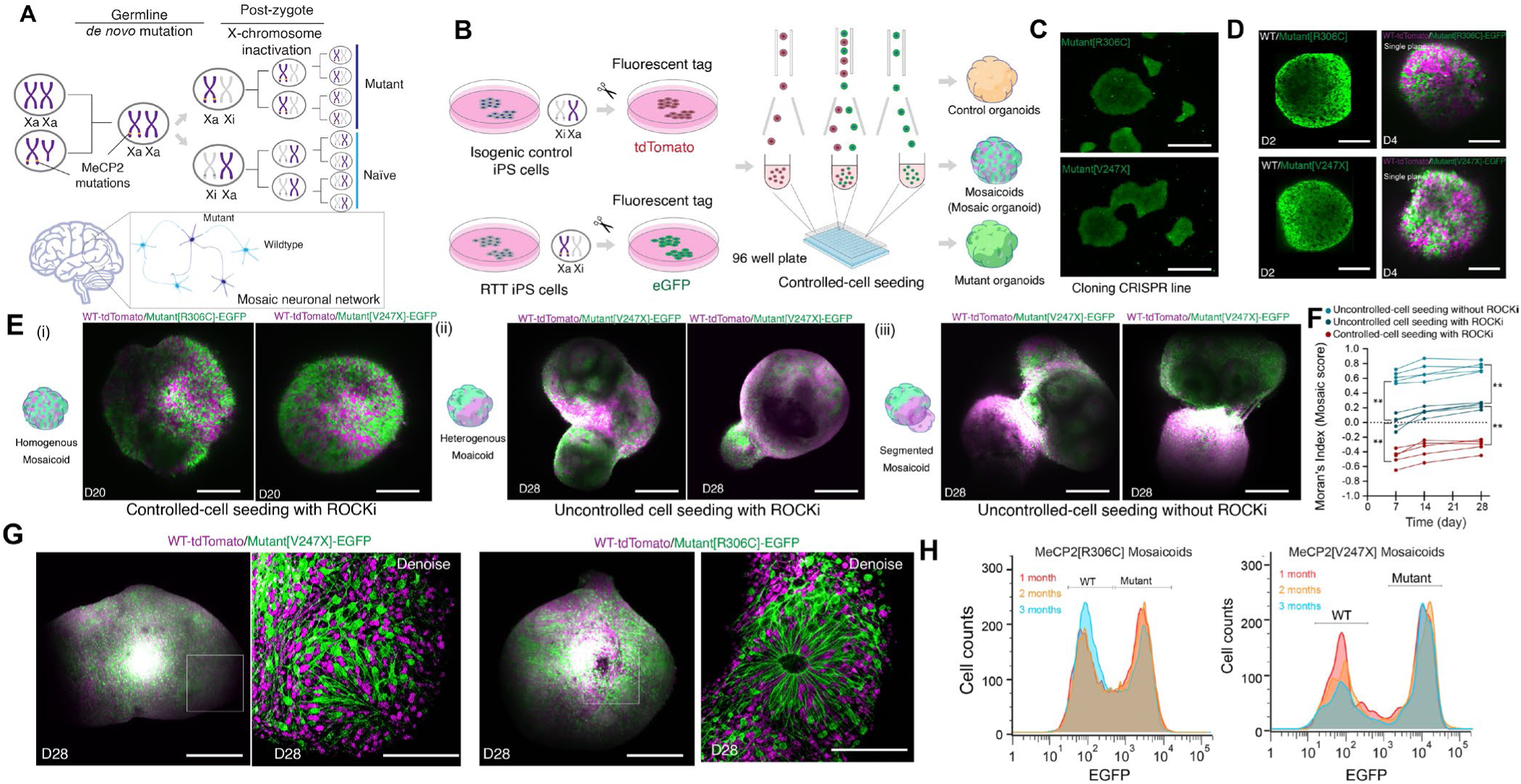
Controlled single-cell seeding generates reproducible human cortical mosaicoids with defined genotype composition. (**A**) Generation of mosaic neural populations through germline *de novo* mutations combined with post-zygotic X-chromosome inactivation (XCI). Germline *de novo* mutations in MECP2 and subsequent XCI result in heterogeneous populations of wildtype and mutant cells that self-organize into mosaic neural networks. (**B**) Experimental workflow for controlled generation of cortical mosaicoids. Human iPS cells from control and RTT patients are transduced with fluorescent tags (eGFP for mutant or tdTomato for wildtype control). Single cells are precisely seeded at defined ratios into 96-well plates in the presence of ROCKi to control initial cell distribution uniformly. (**C**) Cloning of EGFP-expressing iPS cells by CRISPR/Cas9 knock-in into the AAVS1 safe harbor locus. Scale bars, 500 μm. (**D**) Representative two-photon image of mosaicoid development at D2 and D4. Scale bars, 150 μm. (**E**) Three distinct spatial organization phenotypes emerge depending on seeding conditions. (i) homogeneous distribution with controlled-cell seeding and ROCKi treatment at D12 and D18. Scale bars, 150 μm. (ii) heterogeneous mixing pattern with ROCKi treatment but without controlled seeding at D28. Scale bars, 200 μm. (iii) segmented phenotype at D28 showing spatial separation of genotypes. Scale bars, 200 μm. (**F**) Quantitative Moran’s index to represent mosaic score (−1 to 1) at day 7, 17, 28 for three seeding conditions. n = 5 mosaicoids, **p < 0.01; *p < 0.05; N.S., not significant. Error bars represent ±SD. Two-way ANOVA with Tukey-Kramer post-hoc test. (**G**) Controlled-cell seeding and ROCKi treatment demonstrated fine-scale spatial organization of distinct cell populations after 1 month of culture. Scale bars, 200 μm (left) and 80 μm (right), respectively. (**H**) Longitudinal population change in EGFP expressing mutant cell ratios and non-expressing wildtype populations in mosaicoids by flow cytometry, demonstrating cell-autonomous differential growth rates. 3 mosaicoid pools, >10,000 cells in total count.

### Longitudinal volumetric Mosaicoid two-photon genotype-resolved imaging reveals mutation-dependent spatial segregation

Deep and fast imaging of live organoids using standard two-photon microscopy is challenging due to high light scattering and spherical geometry. While two-photon excitation appropriate for our fluorophores (920 and 1040 nm) penetrates the organoid tissue effectively, emission photons (500-600 nm) scatter extensively in the organoid’s lower hemisphere (far side from the objectives), making collection efficiency the primary limiting factor. We addressed this by employing an acousto-optics-based two-photon microscope with integrated upper and lower non-descanned photomultiplier tubes (PMTs; **Fig. 2A**). This dual-PMT configuration with collection optics positioned underneath the mosaicoids maximized photon capture from the organoid interior, enabling visualization of 70-80% of the volume in live organoids and 90-95% in fixed organoids (**Fig. 2B, Fig. S2A**). Combined with refractive index matching using iodixanol^44^, this approach enabled high-fidelity visualization of near-complete organoid volumes, revealing distinct subcellular compartments including surface axonal layers, neuronal soma, neural progenitors, and radial glia populations in deep organoid tissues (**Fig. 2C, Fig. S2B**). Spatial mapping at 3 months of development revealed that mutant and wildtype neurons organize in mutation-type-dependent patterns within mosaicoids (**Fig. 2D, E, Fig. S2C**). Differences in neuronal migration rates and cell-cell adhesion properties generated distinct spatial clustering geometries (**Fig. 2D, E**). V247X mosaicoids exhibited pronounced spatial skewing with mutant-wildtype inhomogeneous segregation (**Fig. 2E, G**) in both radial (R) and angular (theta) cell distributions, whereas R306C mosaicoids displayed clustering against the theta axis (**Fig. 2E, F**). This theta distribution denotes the consistency of emerging cortical columns in human cortical development^23,24^. In addition, we found wildtype neurons tended to be more dominant in superficial layers compare to mutant neurons in V247X mosaicoids (**Fig. 2G**). These mutation-type-dependent spatial phenotypes indicate that distinct MECP2 truncations differentially affect cell proliferation and migration dynamics during mosaicoids development, potentially contributing to genotype-dependent functional alterations observed in RTT. Taken together, it suggests that mosaicism in RTT and X-linked neurodevelopmental disorders manifests not as a stochastic, uniform intermixing of mutant and wildtype cells, but rather as a structured, column-by-column segregation within the developing cortex^25,26^. Because cortical columns represent discrete functional processing units^27^, spatially skewed mutant cell enrichment within individual columns may generate column-specific functional deficits. More generally, the angular bias observed in mosaicoids suggests radial sector-like organization, evocative of developmental patterning, although organoids do not fully recapitulate mature cortical columns.

**Figure 2.**
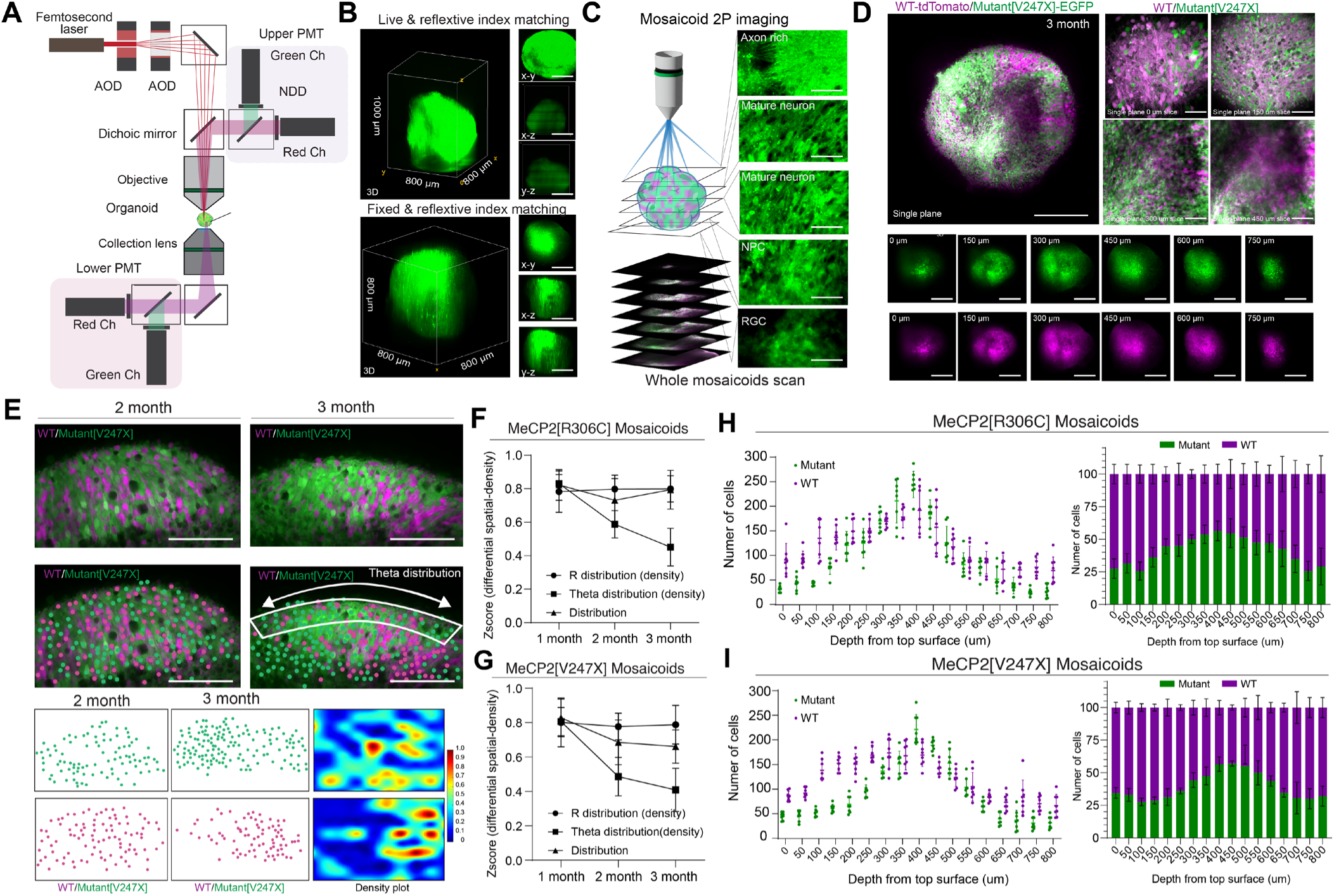
Mutation-dependent spatial organization in developing mosaicoids. (**A**) Schematic of dual-PMT-paired two-photon microscopy with upper and lower PMTs for optimized photon collection throughout the organoid volume. (**B**) Representative images demonstrating enhanced imaging depth with dual-PMT collection (upper + lower PMT) with organoid clearing using iodixanol refractive index matching. Scale bars, 300 μm. (**C**)Mosaicoid two-photon imaging of Mosaicoids represented the unique cellular morphology depending on the imaging depth. Z-stack series reveals distinct neuronal soma, axonal, neural progenitor layers, and RGCs. Scale bars, 50 μm. (**D**) Genotype-resolved (green = wildtype; magenta = mutant) structural imaging of mosaicoids. (**E**) Spatial organization at 2 and 3 months. Representative images of R306C and V247X mosaicoids with quantification of radial (R) and angular (theta) cell distributions. Spatial density maps reveal homogeneous mixing in R306C mosaicoids and clustered segregation in V247X mosaicoids. Scale bars, 200 μm. (**F, G**) Temporal quantification of differential spatial-density Z-scores for R distribution (density), theta distribution (density), and overall distribution in MeCP2[R306C] (**F**) and MeCP2[V247X] (**G**) mosaicoids at 1, 2, and 3 months. V247X mosaicoids exhibited pronounced spatial skewing, with mutant-wildtype inhomogeneous segregation in both theta and R distribution, compared to MeCP2[R306C] mosaicoids. n = 6 mosaicoids, **p < 0.01; *p < 0.05; NS. Error bars represent ±SD. One-way ANOVA with Tukey-Kramer post-hoc test. (**H, I**) Depth-resolved cell distribution analysis of mutant and wildtype cells from the organoid top surface (0-800 μm) in MeCP2[R306C] and MeCP2[V247X] mosaicoids. Proportional composition identified MeCP2[V247X] mosaicoids exhibit pronounced skewed mutant-wildtype segregation across imaging depth, whereas MeCP2[R306C] mosaicoids display a more uniform depth distribution. n = 6 mosaicoids, **p < 0.01; *p < 0.05; N.S., not significant. Error bars represent ±SD. Two-way ANOVA with Tukey-Kramer post-hoc test.

### MosaicoidSim identifies adhesion, growth, and migration parameters sufficient to reproduce mosaic spatial organization

Longitudinal imaging of mosaic organoids generated through controlled cell seeding (**Fig. 1**) revealed complex spatial organization patterns: heterogeneous cell distribution across entire organoids and layer-specific, column-like arrangements within developing cortical domains (**Fig. 2**). To translate these empirical observations into principles governing human cortical mosaic development, we established MosaicoidSim, a computational framework that models organoid growth and cell migration in mosaic contexts. MosaicoidSim systematically interrogates how initial seeding distributions, genotype-dependent cell adhesion^45^, differential proliferation/differentiation rates, and neuronal migration dynamics determine spatial organization and mosaic composition over time (**Fig. 3A, B, see Methods**). Key parameters including the doubling time of cells at each stage and migration rates were determined based on previous literature^46–50^. *In vitro* experiments demonstrated that initial seeding conditions significantly influence whether cells distribute homogeneously or heterogeneously following culture as well as the presence and absence of ROCK inhibitor. Small aggregates formed by uncontrolled seeding resulted in heterogeneous spatial distributions (**Fig. 1D**). *In silico* simulations incorporating varying cell-adhesion parameters successfully recapitulated experimentally observed scenarios, including uniform single-cell seeding (i), uncontrolled seeding with aggregation (ii), and uncontrolled seeding with strong aggregation in the absence of ROCKi, which leads to downregulated cadherin expression^51^, yielding spatial distribution outcomes closely matching those observed *in vitro* (**Fig. 3C**). To assess whether and how the adhesion between wildtype and mutant cells regulates this uniform distribution, we introduced a segmentation index and Moran’s index, and a mosaic score^52,53^ (**Fig. 3D**). Parameter sweeps across the adhesion factors revealed that mosaicoid spatial organization emerged from non-linear interactions between multiple parameters. Small changes in homotypic adhesion coefficients dramatically shifted segregation outcomes from mixed (SI<0.5) to highly segregated (SI>0.8) distributions (**Fig. 3E, Fig. S3**). When WT-Mut adhesion was improved by administration of ROCK inhibitor, regardless of WT-WT adhesion and Mut-Mut adhesion, the mosaicoids achieved uniform state. However, downregulated adhesion between WT-Mut cells in the mosaicoids (in the absence of ROCKi) yielded higher segmentation and ununiform distribution of cells in mosaicoids (**Fig. 3F, Fig. S4**).

**Figure 3.**
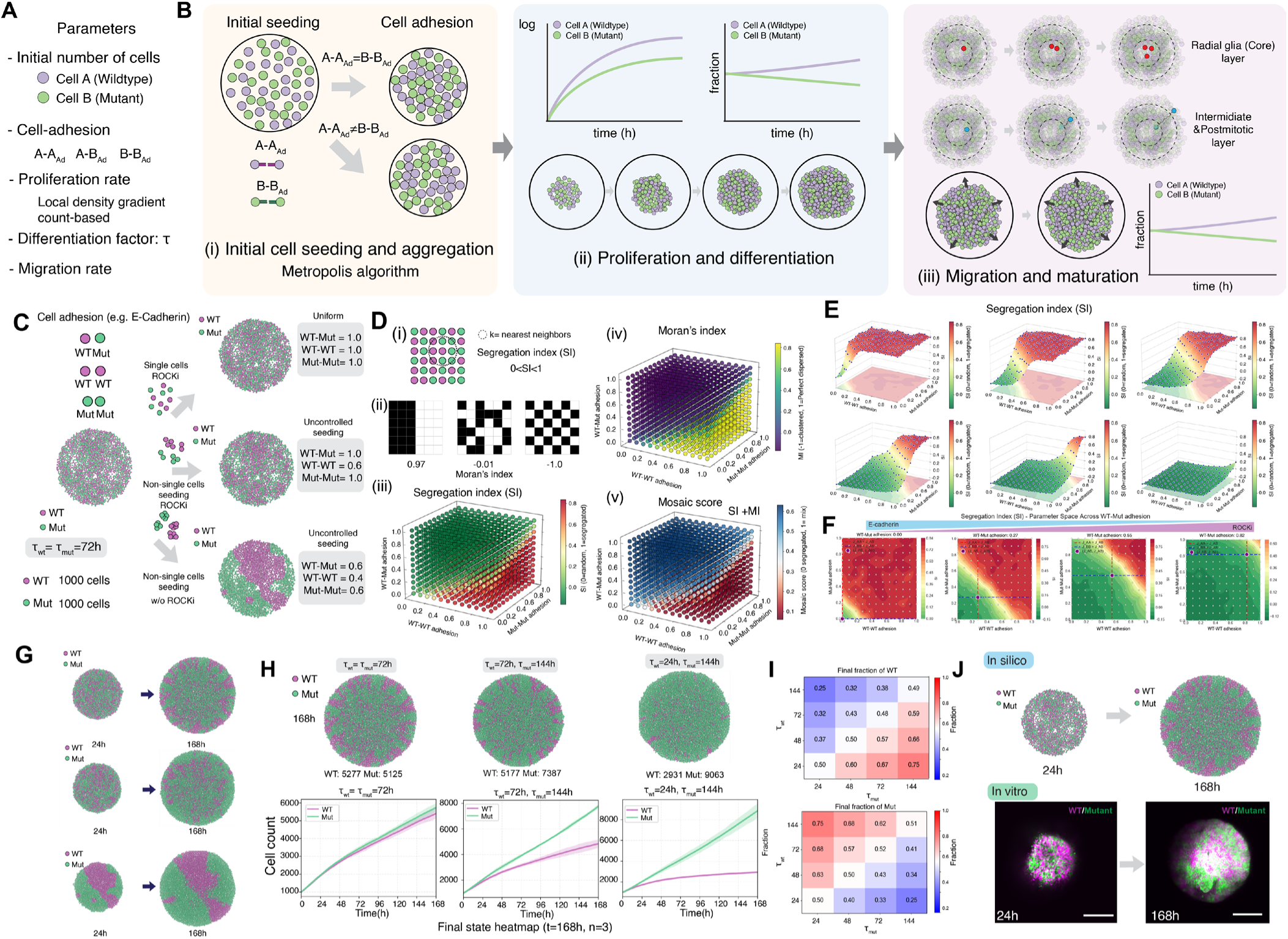
Predicting adhesion-dependent spatial organization in Mosaicoids. (**A**) The computational framework MosaicoidSim incorporates: cell genotype (wildtype and MECP2-mutant), initial cell number (n = 1,000), genotype-specific cell adhesion factors reflecting cadherin expression levels, proliferation restriction parameters, local density gradient count-based measurements, differentiation factor (τ), and genotype-specific migration rates. (**B**) Three-phase developmental model workflow. (1) Initial cell seeding and aggregation via Metropolis-type swap sorting according to adhesion parameters (WT-WT, WT-Mut, Mut-Mut). (2) Proliferation and differentiation. (3) Migration and maturation. Right graphs quantify population composition and spatial segregation over developmental time. (**C**) Adhesion parameter sensitivity analysis. Three representative scenarios with distinct cell-cell adhesion parameters, non-segregating mosaic (uniform mixing), single-zone segregation (modest spatial separation), and unstructured cell (maximal segregation). Corresponding quantitative graphs show population ratios and spatial segregation metrics over time. (**D**) Spatial segregation analysis: (i) Neighbor composition matrices; (ii) Segregation index (SI) calculation; (iii-v) 3D surface plots depicting segregation index, Moran’s index, and mosaic score as functions of adhesion parameters. (**E**) 3D segregation index landscapes across parameter ranges, showing segregation patterns for different genotype combinations. (**F**) 2D heatmaps representing segregation metrics under various adhesion conditions. (**G**) Temporal evolution and quantitative analysis of population dynamics. Cell counts and radius measurements (r_WT/r_Mut; T_WT/T_Mut) indicate genotype-specific spatial localization and temporal changes. (**H**) Final spatial distribution heatmaps at 168 hours. WT organoid (all wildtype cells) showed uniform spatial distribution. Mosaicoids showed depth-dependent spatial organization. (**I**) Heatmaps display final fraction of mutant cells across spatial bins, revealing mutation-type-dependent segregation patterns that correlate with experimental observations. (**J**) Similarity of pattern of organoid growth *in silico* ( τ wt and τ mut =72h) and *in vitro*.

Differences in proliferation rates between wildtype and mutant cells, driven by genotype-dependent differentiation efficiency (τ_WT_ and τ_Mut_), led to skewed growth and unequal cell numbers within mosaicoids (**Fig. 3G**) consistent with *in vitro* observations in V247X mosaicoids (**Fig. 1F**). Systematic sweeping of τ values across wildtype and mutant cells revealed that this factor drives imbalanced cell proportions over time (**Fig. 3H, Fig. S5A-H**). Even under uniform initial seeding conditions, directional migration toward oxygen-rich outer surfaces generated column-like spatial arrangements. These layered structures resembled cortical lamination patterns observed *in vivo*, suggesting that organoid morphology naturally emerges from gradient-driven dynamics without explicit specification of layer formation (**Fig. 3I**). Side-by-side comparison between *in silico* predictions and *in vitro* experimental organoids demonstrated their similarity of spatial organization (**Fig. 3J**). At the 24 hour timepoint, predicted cell distributions closely matched fluorescently-labeled organoid imaging. By 168 hours, both simulations and experiments showed distinct wildtype and mutant spatial domains with layer-specific organization. During neocortical development, MeCP2 expression increases alongside DNA methylation^54^ and MeCP2 has been implicated in progenitor proliferation and neuronal differentiation through Notch signaling^55^. In organoid studies, MECP2-mutant lines exhibit increased proliferation and altered fate determination relative to wild type^56^ and rewire their fate determination. Based on these findings describing cell autonomous effects induced by MECP2, and our simulations that recapitulate mosaicism and demonstrate that cell fates are critical determinants of mosaic spatial outcomes, we suggest that MosaicoidSim captures some fundamental principles governing initial cortical pattern formation in mosaicoids.

We next extended MosaicoidSim to incorporate radial migration in cortical development. Mosaicoids undergo progressive laminar organization during development, with distinct core, intermediate, and peripheral regions emerging over time (**Fig. 4 (i, ii)**). We established a migration model, where each cell migrates toward a layer at a different migration speed for WT and Mut (see **Methods**), to identify how proliferation and migration balance influenced spatial organization after development (**Fig. 4A (iii)**). When we assigned different migration speed parameters, WT and Mut cells exhibited different migration patterns but maintained column-like spatial arrangements (**Fig. 4B, C**). By 288 hours, WT cells were more enriched in superficial layers than Mut cells (**Fig. 4C**), maintaining column-by-column segregation, which is also consistent with *in vitro* results (**Fig. 2E, F, G**). This centrifugal redistribution continued through 408 hours and 672 hours, establishing skewed layer-specific organization. Within *in silico* R306C mosaicoids (50:50 ratio), WT cells migrated faster through core regions and intermediate layers, while mutant cell remained in core and intermediate layers (**Fig. 4D, Fig. S6A, B, C**). This resulted in WT cells preferentially occupying peripheral regions at 672 h, while Mut cells remained concentrated toward core and intermediate regions (**Fig. 4E**). Although we assigned different migration speed and proliferation restriction at postmitotic layers, R306C mosaicoids showed relatively balanced WT and Mut total population (**Fig. 4F**), which is consistent with flow cytometry results (**Fig. 1**).

**Figure 4.**
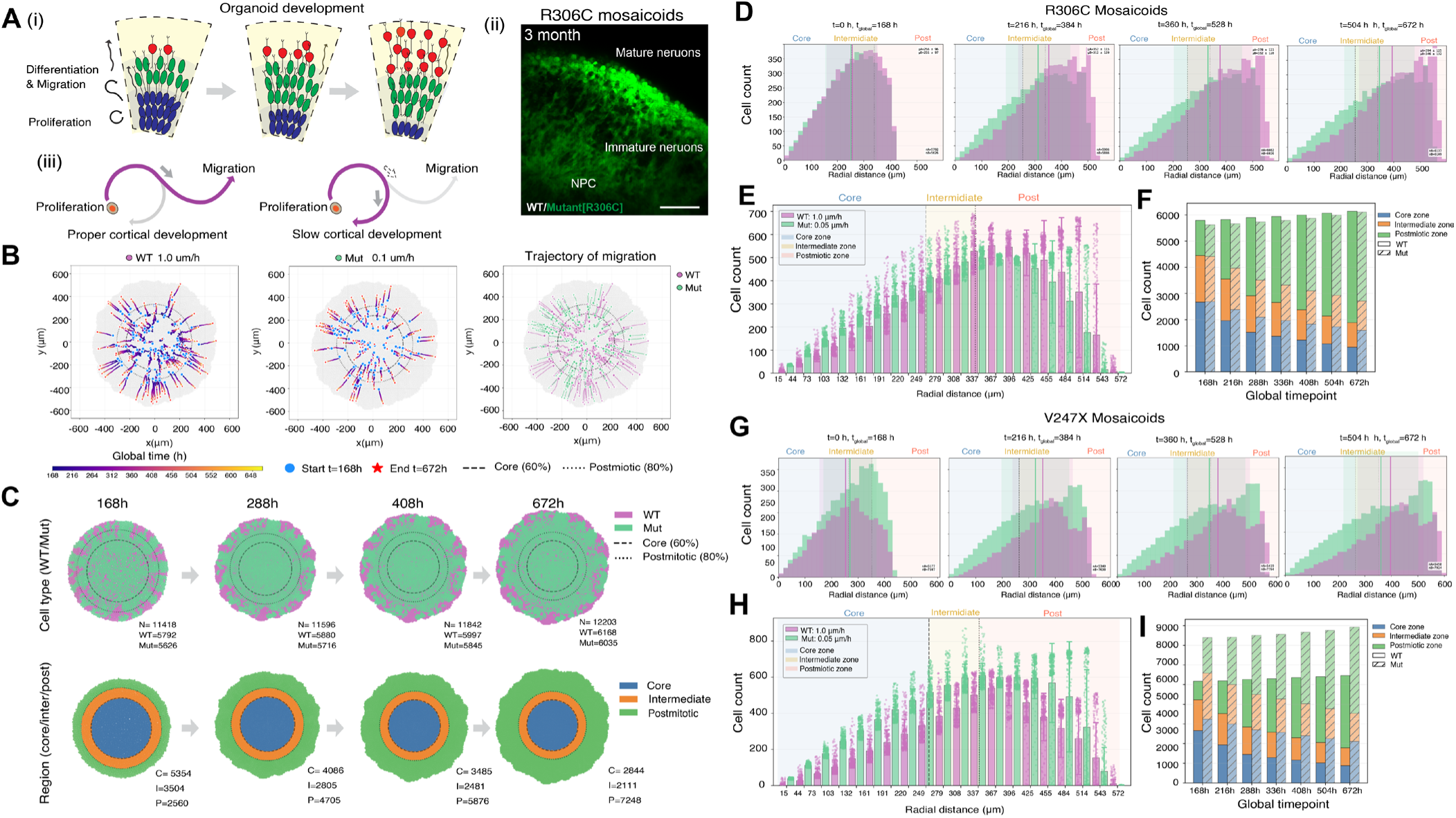
MosaicoidSim validates migration pattern in cortical mosaicoids. **(A**) Schematic of organoid developmental stages showing progressive differentiation and migration (left, i), resulting in layered organization with core (blue), intermediate (green), and mature neuron (red) regions. Representative image of R306C mosaicoids at 3 months showing mature neuronal layer organization (right, ii). Scale bar: 200 μm. (iii) Proliferation–migration balance. Normally, progenitors gradually exit the proliferation cycle and transition to migration (left); delay of transition to migratory neuron may impact this, slowing neocortical development. (**B**) Representative cell migration trajectories (100 cells) for WT and Mut cells at different migration speed, WT: 1.0 μm/h, Mut: 0.1 μm/h. Trajectories emanate from central seeding region toward peripheral regions. (**C**) Longitudinal tracking of cell distribution over developmental time (168h to 672h). Top row shows spatial organization with core (blue, innermost), intermediate (green), and outer (red, periphery) regions. Bottom row shows quantification of cell proportion changes in each ring over time. (**D**) Longitudinal histogram of cell population in R306C mosaicoids. (**E**) Left: Stacked bar chart quantifying total cell counts across spatial regions in R306C mosaicoids, stratified by developmental stage. Right: Cumulative cell count trajectory over time showing total population expansion. (**F**) Schematic illustration of tradeoff relationship between migration and proliferation. Number of bins = 20 μm. Bin width = 30 μm. (**F**) Cell count stratification by spatial region for R306C mosaicoids in core, intermediate, postmitotic layer. (**G, H**) Longitudinal histogram of cell population in V247X mosaicoids showed Mut depletion in superficial layer and excessive accumulation in core and intermediate regions. (**I**) Cell count stratification by spatial region for V247X mosaicoids (right) in core, intermediate, postmitotic layer.

To model V247X mosaicoids, we determined the parameters of different initial cell populations at phase 3 based on flow cytometry results (**Fig. 1**). V247X mosaicoids demonstrated dramatic Mut depletion in superficial layers and excessive accumulation in core and intermediate regions (**Fig. 4G, H and Fig. S7A, B, C**), while preserving the imbalance in total number of cells. This result suggested that initial cell adhesion (phase 1) and proliferation (phase 2), not migration dynamics, is a primary determinant of change of total cell number. Indeed, differential migration dynamics impact the spatial organization of mosaicism (**Figs. 4I**). More generally, the *in silico* findings support the *in vitro* results demonstrating that spatial organization within mosaicoids reflects MECP2 mutation-dependent differences in proliferation, migration, neuronal maturation, and regional cell population dynamics during mosaic cortical development. The spatially skewed enrichment of mutant cells within individual cortical columns and specific layers may generate column- and layer-specific functional deficits in mosaicism.

### Genotype-resolved functional connectivity reveals asymmetric network organization

To investigate how mosaic expression of mutant MeCP2 influences neural circuit assembly during human cortical development in RTT, we developed an integrated platform combining controlled-cell seeding with functional calcium imaging using two-photon microscopy followed by graph theory-based network analysis^19^ (**Fig. 5A**). We identified the genotype in mosaicoids by CRISPR-based fluorescent labeling and characterized neuronal connectivity by functional calcium imaging. In addition, we classified the cell type (inhibitory and excitatory neurons) and cortical layer using AAV-based phenotype identification at single-cell resolution, and pseudo-simultaneous multi-layer calcium imaging, respectively (**Fig. 5B, C**). This allowed us to distinguish spontaneous neural activity in both WT and mutant populations. The corresponding functional connectivity graphs constructed from pairwise activity correlations highlighted distinct genotype-dependent network topologies in both R306C and V247X mosaicoids (**Fig. 5D**). In both R306C and V247X mosaicoids, WT-WT edges exhibited significantly stronger connectivity weights than WT-Mut and Mut-Mut pairs, with no significant difference observed between WT-WT and Mut-Mut connections in the R306C mosaicoids (**Fig. 5F, G**). Quantification of edge counts confirmed that WT-WT pairs formed significantly more connections than WT-Mut and Mut-Mut pairs (**Fig. 5E**, left). This double deficit indicates that MECP2 loss impairs network participation at multiple levels. V247X mosaicoids displayed significantly fewer WT-Mut edges than R306C mosaicoids, suggesting that the V247X truncating mutation produces a more pronounced disruption of cross-genotype connectivity than the R306C mutation (**Fig. 5G and H**).

**Figure 5.**
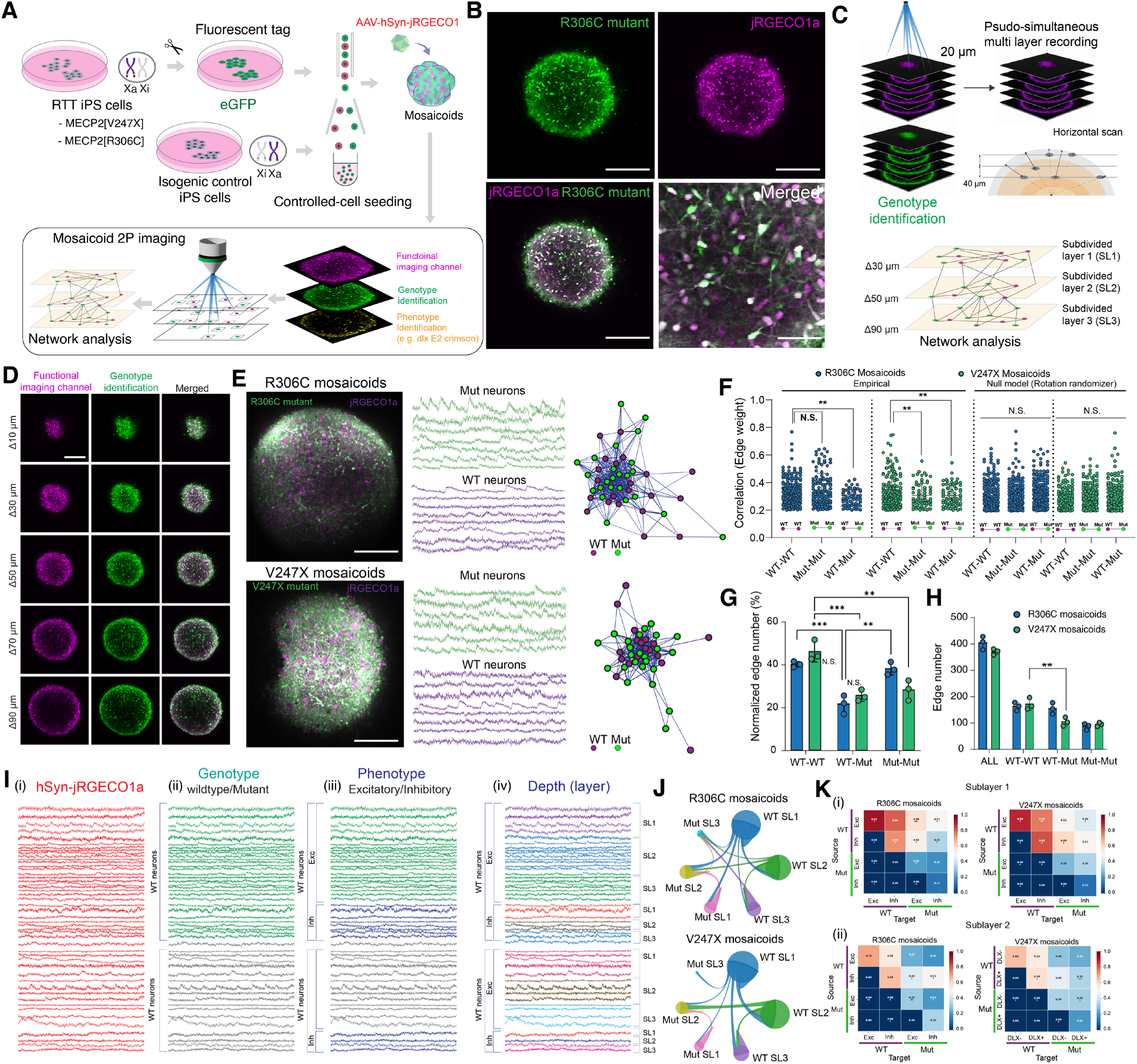
Two-photon calcium imaging of mosaicoids with genotype and phenotype identification revealed imbalance connectivity between WT and mutant neurons. (**A**) Generation of mosaicoids followed by AAV transfection for jRGECO1a. (**B**) Representative R306C and V247X mosaicoids showed jRGECO1a expression (calcium indicator) and genotype-specific labeling (EGFP). Scale bars, 200 μm. (**C, D**) Multi-layer imaging was performed to capture layer-dependent activity. Scale bars, 150 μm (**E**) Calcium recordings and network connectivity maps showing functional connections between wildtype and mutant neurons. Scale bars, 200 μm (**F**) Functional connectivity analysis comparing empirical data versus random null model for different cell-type pairs (WT-WT, Mut-Mut, WT-Mut) in R306C and V247X mosaicoids. n = 6, **p < 0.01; *p < 0.05; N.S., not significant. Error bars represent ±SD. One-way ANOVA with Tukey-Kramer post-hoc test. (**G**) Normalized or (**H**) non-normalized edge numbers demonstrated mutation type-dependent differences in connectivity strength. Edge counts are normalized to the total number of possible pairwise connections within each category. n = 3, **p < 0.01; *p < 0.05; N.S., not significant. Error bars represent ±SD. One-way ANOVA with Tukey-Kramer post-hoc test. (**I**) Calcium imaging traces recorded from neurons expressing (i) raw ΔF/F fluorescence signal with jRGECO1a, (ii) genotype, distinguishing WT from mutant neurons, (iii) phenotype, grouping neurons by cell type, excitatory (Exc) and inhibitory (Inh), and (iv) sublayer (SL1, SL2, SL3). (**J**) Functional connectivity between WT and mutant (Mut) neuron populations across sublayers (SL1, SL2, SL3) in R306C mosaicoids (top) and V247X mosaicoids (bottom). Nodes represent discrete neural populations (e.g., WT SL1, Mut SL2), with node size proportional to total communication number. (**K**) Heatmaps quantifying pairwise functional connectivity strength between neuron populations, stratified by genotype (WT vs. Mut) and cell type (Exc vs. Inh) in (i) Sublayer 1 and (ii) sublayer 2. n = 3 mosaicoids, **p < 0.01; *p < 0.05; N.S., not significant. Error bars represent ±SD. Two-way ANOVA with Tukey-Kramer post-hoc test.

### Sublayer- and cell-type-specific connectivity patterns reveal heterogeneous network vulnerability

To determine whether network deficits are uniform across cortical depth and cell types, or show selective vulnerability in specific populations, we stratified our connectivity analysis along these biological dimensions. The superficial postmitotic zone was subdivided into three bins of equal thickness (sublayers 1–3, from inner to outer, 30µm), and neurons were further classified by phenotype (inhibitory and excitatory neurons) and genotype to enable systematic dissection of connectivity at multiple organizational levels (**Fig. 5I**). To quantify inter-neuronal functional coupling, we constructed pairwise correlation-based connectivity networks across genotype-defined populations (**Fig. 5J**). We found that WT-WT connections were consistently stronger and more numerous than WT-Mut or Mut-Mut connections in both R306C and V247X mosaicoids (**Fig. 5J**). This pattern was evident across all sublayers (SL1–SL3), with WT neurons in upper sublayers (SL1) exhibiting particularly dense intra-genotype connectivity, while mutant neurons formed comparatively sparse inter-population edges. To further characterize connectivity at the cell-type level, we computed pairwise functional connectivity matrices between excitatory and inhibitory subpopulations, stratified by genotype (WT vs. Mut) and cortical layer (**Fig. 5K**). Cre expression under dlx promoter combined with FLEX-E2crimson reporter allow us to resolve its cell type on top of mosaic classification (**Fig. S8**). In sublayer 1, both R306C and V247X mosaicoids showed significantly stronger WT-WT connectivity compared to Mut-Mut pairs across both excitatory and inhibitory subtypes (**Fig. 5K (ii)**). In addition, WT excitatory-to-inhibitory connectivity was preserved while mutant excitatory neurons showed reduced coupling to all target populations, suggesting a selective impairment of Mut-excitatory input/output. In sublayer 2, connectivity patterns diverged further between mutation types (**Fig. 5K (ii)**), but with less correlation compared to sublayer 1, suggesting that interneuron subtype identity partially modulates vulnerability to MECP2 loss-of-function. Taken together, these findings indicate that MECP2 mutations progressively exclude mutant neurons from coordinated network activity and disrupt inter-genotype connectivity in mosaicism. This effect is mutation type-, layer-, and cell type-dependent, arising through multiple cell-autonomous and non-cell-autonomous mechanisms. Distinct mutation classes (R306C and V247X) may thus engage partially non-overlapping pathological pathways despite producing similar network-level deficits.

### Cell-autonomous and non-cell-autonomous transcriptional alterations in mosaicoids

To dissect whether the network abnormalities observed in mosaic organoids arise from cell-autonomous defects, non-cell-autonomous influences from neighboring genotypes, or a combination of both, we performed FACS-based cell isolation of WT and mutant cells from non-mosaic (100:0 or 0:100) organoids and mosaicoids, followed by RT-PCR analysis (**Fig. S9A**). This experimental design enabled comparison of gene expression in cells of the same genotype grown in either pure or mosaic environments (WT vs. WT-Mos; Mutant vs. Mutant-Mos). In both R306C and V247X mosaic organoids, MECP2 expression was reduced in mutant cells compared with WT cells, as expected (**Fig. S9B**). In addition, WT cells grown in mosaicoids (WT-Mos) showed significantly reduced MECP2 expression compared with WT cells in non-mosaic organoids and mutant cells in mosaicoids (Mutant-Mos) showed even further reductions relative to mutant cells in non-mosaic organoids. This indicates that the mosaic environment exerts a non-cell-autonomous influence on gene expression in genotypically WT cells. Examining downstream MECP2 target genes, HDAC2 was significantly upregulated in R306C mutant cells compared with WT cells (**Fig. S9C**). Comparison of mutant cells in non-mosaic versus mosaic conditions revealed that HDAC2 expression was downregulated in mutant cells when surrounded by WT cells in mosaicoids, suggesting partial transcriptional rescue of mutant cells through interactions with WT neighbors in R306C mosaicoids. In V247X mosaicoids, while GABRA2 expression was unchanged between WT and V247X mutant cells, GABRA4 was significantly reduced in mutant cells (**Fig. S9D**). Indeed, impaired excitatory–inhibitory connectivity associated with GABRA4 downregulation, together with reduced protein expression confirmed by immunostaining, was also observed in non-mosaic, fully mutant V247X organoids, consistent with our previous findings^19^. Similarly, when mutant cells were placed in a mosaic environment, both GABRA2 and GABRA4 expression were significantly upregulated compared with non-mosaic cell in mutant organoids (**Fig. S9D**), demonstrating that the mosaic context can partially restore GABAergic gene expression in V247X mutant cells through non-cell-autonomous mechanisms. These results reveal that mosaic MECP2 loss drives both cell-autonomous transcriptional dysregulation and non-cell-autonomous remodeling, with bidirectional cross-talk. WT cells shift partially toward a mutant-like state, while mutant cells partially recover toward WT ^57^.

### MosaicoidSeq identifies cell-type-specific transcriptional alterations and predicted synaptic communication deficits

In parallel with functional imaging of mosaicoids, we employed single-cell RNAseq with multiplex genotype annotation as MosaicoidSeq to systematically distinguish cell-autonomous and non-cell-autonomous effects of MECP2 mutations at single-cell resolution (**Fig 6A**). This demultiplexing strategy leveraged two orthogonal demultiplexing signals, mutation-specific SNVs identified by WES and EGFP read status, to unambiguously assign WT or Mutant identity to each transcriptome within shared t-SNE clusters (**Fig 6A**). This dual-anchor demultiplexing strategy achieved robust genotype resolution and enabled direct, same-environment transcriptomic comparisons between WT and mutant cells in mosaicoids, providing a powerful platform for investigating MECP2 mutation-dependent transcriptional and communication phenotypes. Integrated t-SNE analysis of MosaicoidSeq resolved distinct cell type populations (**Fig. 6B**), encompassing the full spectrum of cortical organoid cell types. Genotype-stratified analysis revealed that MECP2 R306C and V247X mutations produced distinct cell population distributions compared to WT cells (**Fig. 6C, D**). V247X mutant cells gave rise to a transcriptomically unique excitatory neuron cluster that was enriched in the mutant condition and absent in WT counterparts (**Fig. 6D**). Cluster identities were validated by canonical marker expression: VIM and SOX2 delineated radial glia and progenitor populations; DCX and NEUROD6 marked excitatory neuronal lineages; ASCL1 and GAD2 identified inhibitory lineages; GRIA2 and GRIN2B confirmed glutamatergic identity; and APOE, S100B, SATB2 annotated astrocyte and GABAergic neuron populations, respectively (**Fig. 6E, F**).

**Figure 6.**
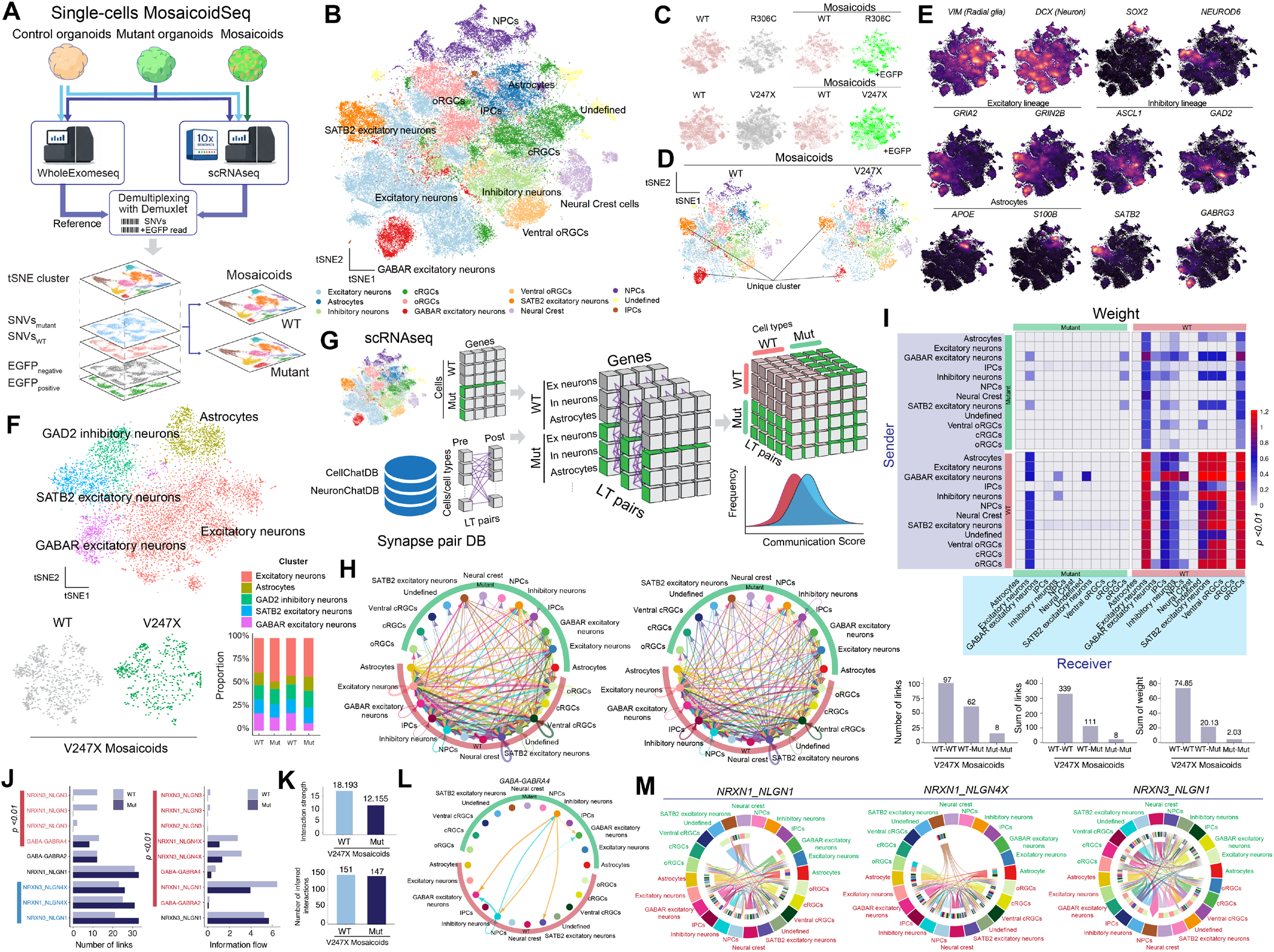
MosaicoidSeq reveals cell-type-specific transcriptional disruptions and attenuated intercellular communication networks in mosaicoids. (**A**) MosaicoidSeq pipeline. MECP2-mutant (EGFP-labeled) and wild-type (WT) cortical organoids and Mosaicoids were prepared, and dissociated single cells were subjected to parallel whole-exome sequencing (WES) and scRNA-seq. Genotype identities (WT vs. mutant; EGFP⁻ vs. EGFP⁺) were resolved post hoc, leveraging mutation-specific single-nucleotide variants (SNVs) and EGFP reads as orthogonal demultiplexing anchors. (**B**) Integrated t-SNE visualization of all single cells from Mosaicoid datasets. (**C**) t-SNE plots illustrates the genotype-specific distribution of transcriptomic populations across the shared embedding. (**D**) Transcriptomically unique excitatory neuron clusters selectively loss in the V247X mosaicoids. (**E**) Canonical cell-type markers across the integrated t-SNE. (**F**) t-SNE visualization of V247X Mosaicoids highlighting GAD2⁺ inhibitory neurons, SATB2⁺ excitatory neurons, unique excitatory neurons, astrocytes, and excitatory neurons. (**G**) Inferring intercellular communication pipeline using NeuronChatDB (Synapse Pair DB) to infer ligand-receptor (LT) interaction pairs. Communication scores were quantified and compared between WT and mutant cells across all cell-type sender-receiver matrices. (**H**) Global cell-cell communication networks in weight (left) and number (right) of links in mosaicoids. Line thickness and color denote the strength and origin of intercellular interactions. (**I**) Heatmap of differential communication weights across all sender-receiver cell-type pairs in mutant versus WT conditions. Bottom showed the number of links, sum of links, and sum of weights for WT-WT, WT-Mut, and Mut-Mut interactions. (**J**) The number of significant links for specific LT pairs differentially regulated between WT (dark) and mutant. (**K**) Total information flow and the number of altered interaction pairs in V247X Mosaicoids, indicating a global reduction in intercellular signaling strength. (**L**) GABA-GABRA4 signaling pathway interactions across cell types in V247X Mosaicoids, demonstrating widespread GABAergic communication originating. (**M**) Chord diagrams for three individual trans-synaptic Neurexin-Neuroligin ligand-receptor pairs-NRXN1_NLGN1, NRXN1_NLGN4X, and NRXN3_NLGN1, illustrating the cell-type-specific distribution and genotype-dependent redistribution of synapse-organizing signaling in V247X mosaicoids.

In sub-cluster analysis, we found that V247X mosaicoids exhibited a significant reduction of this GABA receptor expressing excitatory neuron subtype, accompanied by altered inhibitory excitatory neuron ratios (**Fig. 6F**). These findings indicate that MECP2[V247X] mutation disrupts the fidelity of neuronal differentiation trajectories in a cell-autonomous manner, producing aberrant neuronal subtype identities not observed under WT conditions. Furthermore, to predict whether MECP2 mutations reshape the intercellular signaling landscape beyond cell-intrinsic transcriptional changes, we implemented a comprehensive ligand-receptor (LT) interaction analysis pipeline integrating NeuronChatDB (**Fig. 6G**). Chord diagram visualization of global communication networks inferred altered intercellular connectivity in V247X mutant cells relative to WT, characterized by significantly reduced interaction density across neuronal and non-neuronal cell-type pairs (**Fig. 6H**). Quantitative comparison revealed a dramatic reduction in overall communication weight, with the sum of interaction weights collapsing from 74.85 in WT to 2.03 in V247X mutant cells, a >35-fold decrease, demonstrating a profound and widespread attenuation of intercellular signaling (**Fig. 6I**). Whereas V247X mosaicoids showed marked connectivity asymmetry between mutant and WT cells, R306C mosaicoids showed no comparable imbalance. Inferred interactions were broadly similar in character across WT–WT, WT–Mut, and Mut–Mut pairs, although Mut–Mut pairs consistently carried the highest summed interaction weight and the greatest number of links (**Fig. S10A, B, C**). Rather than the near-complete signaling collapse seen in V247X, R306C mutant cells therefore remain fully integrated into the communication network and are modestly over-connected to one another. This graded difference suggests that the truncating V247X allele disrupts intercellular signaling far more severely than the R306C missense allele, with R306C producing a mild shift in the distribution of interactions rather than a loss of them.

Further analysis of total information flow confirmed this global communication deficit in V247X mutant Mosaicoids (**Fig. 6K**). To identify the specific signaling systems most disrupted in MECP2[V247X] mosaicoids, we interrogated differentially regulated LT pairs (**Fig. 6J**). There was significant reduction of extensive network of GABAergic communication via GABA_GABRA4 emanating from wildtype-inhibitory neuron to multiple target cell types within the Mosaicoids, including mutant excitatory neurons, which is consistent with a previous study ^58^ (**Fig. 6L**). In addition, we identified a distinct sender-receiver cell-type connectivity architecture that specifies how synaptogenic signals are precisely distributed across the organoid cellular network through Neurexin-Neuroligin (NRXN-NLGN) ligand-receptor pairs, including NRXN1_NLGN1, NRXN1_NLGN4X, and NRXN3_NLGN1(**Fig. 6M**). These findings suggest that the MECP2[V247X] mutation reduces predicted GABAergic and NRXN–NLGN-mediated synaptic communication pathways^59^, providing a molecular correlate of the functional connectivity deficits observed by calcium imaging and a molecular and circuit-level basis for the synaptic imbalance and connectivity deficits that define the pathology underlying Rett syndrome mosaicism.

## Discussion

We establish MosaicoidX, an integrated human mosaic organoid platform for engineering, functional imaging, modeling, and molecularly dissecting cellular mosaicism in Rett syndrome. Using isogenic wildtype and MECP2-mutant hiPS cells carrying either R306C or V247X mutations, we generated cortical mosaicoids with prospectively controlled genotype composition by FACS-based single-cell seeding. Key findings establish that mosaicism is not merely a passive mixture of wildtype and mutant cells, but rather an active developmental process that fundamentally shapes tissue architecture, circuit assembly, and intercellular communication.

This approach enabled reproducible formation of homogeneous mosaic organoids when combined with ROCK inhibitor treatment, whereas uncontrolled seeding produced heterogeneous clustering or genotype-dependent segmentation (**Fig. 1**). A major advance of this study is the systematic reconstruction of X-linked mosaicism in a controllable human cortical developmental model. Rett syndrome primarily affects females, in whom random X-chromosome inactivation generates a brain composed of cells expressing either wildtype or mutant MeCP2. However, conventional approaches including patient-derived organoids, non-mosaic mutant organoids^19,60^, animal models^61^, and clinical studies^62^ conflate multiple variables, making it difficult to isolate the effects of random XCI, genotype ratio, spatial organization, and wildtype–mutant interaction. MosaicoidX overcomes this limitation by allowing defined proportions of mutant and wildtype cells to be mixed from the beginning of organoid formation while preserving genotype identity through stable fluorescent labeling. MosaicoidSim recapitulated these patterns and demonstrated that small variations in cell-cell adhesion, proliferation, differentiation, and migration can shift mosaicoids between mixed and segregated states. These findings reveal that tissue architecture in mosaicoids is shaped by both genetic perturbations and biophysical parameters such as cell-cell interactions, initial geometry, and starting population composition. Therefore, standardizing seeding protocols is essential for accurate interpretation of mosaic organoid phenotypes. To our knowledge, this represents the first systematic platform to engineer human cortical mosaicism and analyze its consequences across anatomy, neuronal activity, computational simulation, and single-cell molecular biology.

Longitudinal imaging revealed that mutant and wildtype cells develop mutation-dependent spatial organization, including radial, angular, and layer-associated segregation rather than remaining randomly intermixed (**Fig. 2**). A key finding is that MECP2 mutations autonomously affect cortical development through distinct effects on cell proliferation, adhesion, differentiation, and migration within mosaic situation. In neocortex development, MeCP2 expression progressively increased along with the methylation^54^ and MeCP2 play a role of proliferation and its differentiation involving the notch signaling pathway^55^. In organoid study, MECP2-mutant lines proliferate faster than WT^56^ and rewire their fate determination, which is consistent with our results. The emergence of angular or column-like genotype organization is particularly important, because cortical columns represent local functional processing units^25–27^. Thus, in the mosaic RTT brain, regional enrichment of mutant cells within specific cortical domains may generate local circuit dysfunction, potentially explaining why individuals with similar MECP2 mutations or balanced XCI ratios can show highly variable clinical phenotypes^18^^17^, though further validation with human anatomical assay may be needed. Functional calcium imaging further showed a significant deficit in cross-genotype neuronal communications (**Fig. 5**). WT-WT pairs showed substantially stronger and more frequent functional connections compared to WT-Mut pairs, which in turn exceeded Mut-Mut interactions across both mutations. This indicates that MECP2-mutant neurons are not merely abnormal as isolated cells, but are progressively excluded from coordinated network dynamics. The reduction in cross-genotype connectivity was especially severe in V247X mosaicoids. Sublayer- and cell-type-resolved analyses further showed that these connectivity defects depend on cortical depth and neuronal subtype, with mutant excitatory neurons displaying particularly reduced coupling to other populations^63^. These findings provide direct evidence that mosaic MECP2 mutations disrupt neural circuit assembly through asymmetric wildtype-mutant communication^64,65^.

Finally, single-cell transcriptomic and ligand-receptor analyses revealed impaired intercellular signaling in GABAergic and neurexin–neuroligin communication pathways, particularly involving GABAergic signaling and NRXN-NLGN adhesion pathways^66^ (**Fig. 6**). These pathways are central to inhibitory balance, synapse formation, and network maturation^58^, and their disruption provides a molecular explanation for the impaired functional connectivity observed by calcium imaging. Therefore, the combined imaging and transcriptomic data support a model in which MECP2 mutations impair both intrinsic neuronal maturation and the ability of mutant neurons to communicate with neighboring wildtype cells. Importantly, our data reveal bidirectional non-cell-autonomous effects^67^ of MECP2 mosaicism (**Fig. S9**). Wildtype cells in mosaic organoids showed altered gene expression compared to non-mosaic controls, while mutant cells displayed partial rescue when surrounded by wildtype neighbors, indicating bidirectional transcriptional cross-talk. These findings demonstrate that MECP2 deficiency spreads to neighboring wildtype cells, a process that appears distinct from gap junction-mediated spreading observed in astrocytes^68^. This spreading likely occurs through paracrine signaling between neurons or across neuron-astrocyte interfaces, involving the impaired communication pathways identified here, including disrupted GABAergic and neurexin-neuroligin signaling. Thus, MECP2 mosaicism creates bidirectional interactions in which mutant cells propagate their defects to wildtype neighbors through indirect secreted or signaling-mediated mechanisms, while wildtype cells simultaneously provide partial buffering of mutant dysfunction^57^.

Beyond MeCP2 mosaicism in Rett syndrome, MosaicoidX provides a generalizable framework for studying mosaicism arising from *de novo* mutations and inherited variants. Many X-linked neurodevelopmental disorders, including those involving FMR1, CDKL5, PCDH19, CASK, ARX, and related genes, may be shaped by XCI-dependent cellular mosaicism^69^. Similarly, somatic mosaic mutations affecting pathways such as mTOR, PTEN, TSC1/2, DEPDC5^41^., and other neurodevelopmental regulators can generate focal or distributed mutant cell populations in the brain^70^. In these disorders, disease severity likely depends not only on the presence of mutant cells, but also on their abundance, spatial distribution, developmental timing, cell-type identity, and communication with wildtype neighbors. MosaicoidX enables precise control of these variables to define mosaic threshold effects, identify vulnerable cell populations, and test approaches that restore wildtype–mutant cellular communication.

An important trade-off in our approach is that we engineer mosaic composition by combining isogenic lines, thereby controlling the outcome of XCI without modeling the process itself. This design sacrifices the ability to study how XCI ratios are naturally established and maintained. These include how XCI ratios are established during early embryogenesis, how skewing arises and is selected for, and how the epigenetic remodeling and dosage compensation accompanying XCI unfold during neural lineage specification. Instead, we gain precise prospective control over genotype composition, enabling systematic analysis of mosaic consequences. Addressing these questions will require naïve and re-primed hiPS cell states, in which random XCI can be initiated *de novo* and tracked through differentiation^71–74^. Such complementary systems recapitulate the biological process of XCI but sacrifice the prospective control over composition that the current approach provides. In addition, while two-photon imaging enables genotype-resolved calcium imaging in live mosaicoids, subdividing neurons by multiple categories (e.g. genotype, cortical depth, and excitatory versus inhibitory identity^75^) limits statistical power for some analyses. Future improvements, including larger field-of-view imaging^76,77^, faster multi-plane acquisition, adaptive optics^78^, and improved calcium indicators^79^, will enable analysis of larger neuronal populations while retaining single-cell resolution.

In summary, MosaicoidX establishes a systematic platform for engineering and dissecting human cortical mosaicism in a controllable developmental context, overcoming major limitations of conventional patient-derived stem cell and animal models. Applied to Rett syndrome, this work establishes that MECP2 mosaicism is fundamentally not a static mixture of wildtype and mutant cells, but rather an active developmental mechanism that shapes tissue architecture through mutation-dependent spatial organization, altered proliferation and migration, and progressive exclusion of mutant neurons from developing circuits. At the molecular level, impaired GABAergic and neurexin–neuroligin signaling explains the asymmetric functional connectivity deficits observed by imaging, at least in severe Rett mutations. Critically, our findings demonstrate bidirectional non-cell-autonomous effects, wherein mutant cells perturb wildtype transcriptional states while wildtype neighbors provide partial rescue, a mechanism that likely explains phenotypic variability in Rett syndrome. More broadly, MosaicoidX provides a generalizable framework for modeling genetic mosaicism caused by X-linked and somatic mutations, enabling future studies to define mosaic threshold effects, identify vulnerable cell populations and developmental windows, and test therapeutic interventions that restore wildtype–mutant cellular communication.

## Methods

### Ethics approval

All methods were performed in accordance with the relevant guidelines and regulations. The use of human iPS cells was reviewed and approved by Environmental health safety at Massachusetts Institute of Technology through STR-based human cell-line authentication (MIT). The human iPS cells were handled in accordance with approved protocols.

### Preparation of Rett syndrome patient-derived iPS cells and isogenic controls

Rett syndrome patient-derived hiPS cells carrying the heterozygous MECP2[V247X] mutation (female, 25Y, 705G deletion, frameshift; very rare in RTT patients) and its isogenic control were obtained from the Coriell Institute (WIC07i-07982-4(MT) and WIC07i-07982-2(WT)). The isogenic control cell line had both wildtype and mutant alleles of MECP2, but the mutant allele that had the MECP2 mutation was X-inactivated in all the mutant cells (Fig. S1A). Another Rett syndrome patient-derived hiPS cell line, which carries the heterozygous MECP2[R306C] mutation (female, 8Y, missense; 4-7% of RTT patients), and its isogenic control was obtained from the Coriell Institute (WIC05i-127-325(MT) and WIC04i-127-33(WT)). The isogenic control was established by site-directed genome editing by CRISPR/Cas9 to repair the mutant allele of MECP2, resulting in isogenic control cells having only wild-type alleles of MECP2 (**Fig. S1A**) ^80^. Healthy human iPS cells were also obtained from Coriell Institute (GM23279, female, 36Y) as a non-RTT related healthy control. All hiPS cells were maintained on Matrigel-coated 6-well plates in mTeSR Plus medium (STEMCELL Technologies) with 10 μM Y-27632 (Rock inhibiter, STEMCELL Technologies) for the first day after each passage and sub-cultured every 5-7 days using ReLeSR reagent (STEMCELL Technologies). Cells pluripotency and genotypes were characterized by immunostaining with pluripotent markers (OCT3/4, Nanog, TRA-1-60, and SSEA4), and, allele-specific PCR, whole exome sequencing was performed, that previously described ^19^. All key resources were shown in **Table S1**.

### Generation of fluorescent reporter hiPS cell lines by CRISPR/Cas9 knock-in into AAVS1

To enable genotype-resolved tracking of wild-type and MECP2-mutant cells in mosaicoids, fluorescent reporter cassettes were introduced into the AAVS1 safe-harbor locus of human induced pluripotent stem cells (hiPS cells) using CRISPR/Cas9-mediated homology-directed repair. Reporter donor constructs contain either an EGFP or tdTomato expression cassette flanked by left and right AAVS1 homology arms with AAVS1-Pur-CAG-EGFP (Addgene, #80945) and AAVS1-tdTomato (Addgene, #159275). For genome editing, hiPS cells were dissociated into single cells using TrypLE Express and plated on Matrigel-coated plates in mTeSR Plus medium supplemented with 10 µM Y-27632 ROCK inhibitor 24 h before transfection then detached using TrypLE Express for 5 min. A total of 2×10^6^ human iPS cells were co-transfected with 8 µg of PX458-AAVS1 (Addgene, 113194) plasmid, which expresses SpCas9, and 6 µg of AAVS1-Pur-CAG-EGFP or AAVS1-tdTomato in 100 µl of Opti-MEM. Electroporation was performed using a BTX ECM830 electroporator (Poring plus: voltage, 135 V; pulse length, 5 ms; pulse, 100 ms; number of pulses, 2. Transfer pulse: voltage, 20 V; pulse length, 100 ms; pulse, 100 ms; number of pulses, 5). After electroporation, iPS cells were seeded onto Matrigel-coated plates in mTeSR plus with CloneR2, and 24 hours post-electroporation, transfected iPS cells were subjected to the puromycin selection at 0.25 μg/ml with 10 μM of Y-27632. Then, the culture medium was replaced with mTeSR plus and allowed to recover for one week. Once the cells reached 80% confluence, they were dissociated by treatment with TrypLE Express (Thermo Fisher Scientific) for 5 min. Next, the cell suspension was filtered using 70 µm cell strainers (BD Falcon) in mTeSR plus medium. EGFP or tdTomato-positive cells were sorted using a cell sorter (BD Melody) and re-plated as a single clone to a Matrigel-coated plate in mTeSR plus with CloneR2 to obtain a single colony. Expanded single colonies were then collected for genotyping to confirm targeted insertion and to exclude random integration where applicable. Reporter expression was confirmed by fluorescence microscopy and flow cytometry. Only clones showing stable, uniform fluorescence expression and normal human iPSC morphology were used for mosaicoid generation.

### Controlled cell seeding by cell sorter to form mosaicoids

To generate uniform mosaicoids, fluorescently labeled human induced pluripotent stem cells (hiPSCs) were seeded using fluorescence-activated cell sorting (FACS, BDMelody). MECP2-mutant and corresponding isogenic control hiPSCs carrying genotype-specific fluorescent reporters were maintained on Matrigel-coated plates in mTeSR Plus medium. For mosaicoid generation, hiPSCs were dissociated into single-cell suspensions using TrypLE Express (Thermo Fisher Scientific) for 10–15 min at 37 °C. Dissociated cells were gently triturated, diluted in sorting buffer consisting of phosphate-buffered saline (PBS) supplemented with 1% bovine serum albumin (BSA) and 10 µM Y-27632 ROCK inhibitor, and passed through a 70-µm cell strainer to remove aggregates then, the two cell suspensions were then combined and loaded onto the cell sorter.

Then, this mixed single-cell suspension was sorted using a BD FACSMelody cell sorter. Cells were sequentially gated to exclude debris based on forward and side scatter, to remove doublets using FSC-A/FSC-H and SSC-A/SSC-H parameters, and to isolate viable single cells. Genotype-defined populations were then isolated based on reporter expression: EGFP-positive cells and EGFP-negative were collected as separate populations. For controlled mosaicoid formation, WT and MECP2-mutant hiPSCs were directly sorted into U-bottom ultra-low attachment 96-well plates (Corning) containing mTeSR Plus supplemented with 20 µM Y-27632. Unless otherwise indicated, a total of 30,000 cells were seeded per well at 1,000 event/sec sorting speed. Mosaicoids were generated at predefined WT:mutant ratios, including 50:50 as appropriate. For direct FACS-based seeding, the sorter was programmed to deposit the required number of cells from each genotype into each well. In parallel experiments, sorted cells were collected separately, counted, mixed at defined ratios, and plated at the same total cell number. After 24 h, the medium was replaced with neural induction medium to initiate cortical differentiation. Mosaicoids were subsequently differentiated using the same cortical organoid protocol as non-mosaic organoids. ROCK inhibitor was maintained only during the initial aggregation period unless otherwise specified. For comparison with controlled FACS-based seeding, uncontrolled mosaicoids were generated by manually mixing dissociated WT and MECP2-mutant hiPSCs at the same nominal ratios without FACS-based singlet deposition. These uncontrolled cultures were established either in the presence or absence of ROCK inhibitor to assess the effect of initial cell distribution and cell adhesion on mosaic tissue organization. After 24 h, the culture medium was replaced with neural induction medium (knockout DMEM-F12, 15% (v/v) knockout serum replacement, 1% (v/v) MEM-NEAA, 1% (v/v) GlutaMAX, 100 nM LDN-193189, 10 μM SB431542, and 10 μM XAV939) and changed every other day. After 10 days of culture, the culture medium was replaced with 1:1 mixture of knockout DMEM/F12 and neurobasal medium supplemented with 0.5% (v/v) N2 supplement, 1% (v/v) B27 plus supplement with vitamin A, 1% (v/v) GlutaMAX, 1% (v/v) MEM-NEAA, and 2.5 ng/ml (v/v) human insulin solution. This medium was changed every 2 days until 18 days of culture had elapsed. After 18 days, the organoids were transferred to flat-bottom ultra-low attachment 24-well plates (Corning) on a rocking shaker (120 rpm) and switched to a maintenance medium (Neurobasal medium supplemented with 1% (v/v) N2 supplement, 2% (v/v) B27 supplement with vitamin A, 1% (v/v) GlutaMAX, 1% (v/v) MEM-NEAA, 250 ng/ml (v/v) human insulin solution, 20 ng/ml BDNF, 200 μM ascorbic acid, 100 μM dibutyryl-cAMP (Sigma-Aldrich), and 1% (v/v) Penicillin/Streptomycin). After 30 days from differentiation, the culture medium was changed twice per week with BrainPhys medium supplemented with 2% NeuroCult™ SM1 Neuronal Supplement, 1% (v/v) GlutaMAX, 20 ng/ml BDNF, 20 ng/ml GDNF, and 1%(v/v) Penicillin/Streptomycin until the time of imaging.

To quantify genotype composition over time, mosaicoids were collected at indicated developmental stages and dissociated into single cells using TrypLE Express. Dissociated cells were filtered through a 70-µm strainer and analyzed by flow cytometry using the same fluorescence gates established during sorting. The relative abundance of WT and MECP2-mutant cells was calculated from the proportion of tdTomato-positive and EGFP-positive viable singlets. Flow cytometry data were analyzed using FlowJo software.

### Virus infection

To visualize neuronal activity by fluorescent Ca^2+^ indicators, pAAV.CAG.GCaMP6f.WPRE.SV40 was a gift from Douglas Kim & GENIE Project (Addgene #100836). pAAV.Syn.NES-jRGECO1a.WPRE.SV40 was a gift from Douglas Kim & GENIE Project (Addgene plasmid # 100854; http://n2t.net/addgene:100854; RRID:Addgene_100854), and AAV-dlx-Cre. Additionally, 2 μL of AAV solution was added to 500 μL of medium in 24-well plates containing one cortical organoid and mosaicoids at least 1-2 weeks before imaging.

### Genotype-resolved two-photon structural and calcium imaging and functional connectivity analysis

For these, we used a two-photon microscope employing a AO-based 3D volumetric imaging (Femtonics3D Atlas) with Insight X3 laser (Spectra-Physics) at 920 nm to excite EGFP/GCaMP6f at Green channels (Semrock, 540/50 nm) and tdTomato at red Channels (Semrock, 610/45 nm) or 1040 nm for jRGECO1a at red Channels (Semrock, 610/45 nm), and 820 nm for E2-Crimson at far red channel (Thorlabs, DMLP650R, long-pass dichroic mirrors: 650 nm cut-on). Functional imaging and single cells-targeting stimulation (ablation) were performed with a 3D acousto-optical multi-photon laser-scanning microscope (Femto3D Atlas, Femtonics). Deflector with 4 acousto-optical crystals allow us to perform high-speed functional imaging and random-access multiplane stimulation with high spatiotemporal resolution (Single cell). Single-plane functional imaging was performed at up to 30 Hz with a 500 × 500 pixel field of view. For multi-plane volumetric functional imaging, three axial planes separated by 30 µm were acquired simultaneously at 10 Hz per plane (500 × 500 pixels each), allowing concurrent recording of activity across the three subdivided layer (SL1-3). For anatomical reference, volumetric z-stacks were acquired with a 2-µm axial step size over a total depth of 600–800 µm. The microscope and laser were controlled MEC software (Femtonics). The dual-PMT configuration, with collection optics positioned above and below the mosaicoids, was used to improve fluorescence collection efficiency. Analog signals from the upper and lower PMTs were merged and routed to the digitizer for synchronized acquisition and downstream analysis. For live imaging, samples were prepared using iodixanol-based refractive index matching to reduce optical scattering and improve fluorescence signal detection during deep two-photon imaging. Briefly, tissues were equilibrated in iodixanol refractive index matching solutions before imaging. The microscope and laser were controlled using MES software/Femtonics control software.

### Data processing of calcium imaging data set

Time-lapse calcium imaging data were processed using Suite2p (Python 3.9) to identify active neurons and define regions of interest (ROIs) encompassing individual neurons within the field of view. For each ROI, baseline fluorescence (F₀) was calculated as the average of the bottom 10th percentile of fluorescence intensity values. The relative change in fluorescence (ΔF/F₀) was computed as (F − F₀) / F₀. Photobleaching correction was applied by normalizing ΔF/F₀ traces using linear regression parameters derived from initial (F_start) and final (F_end) fluorescence values across each time series. For experiments involving mosaicoids with DLX co-labeling, Suite2p was used to identify and export binary masks for two neuronal populations: (1) EGFP-expressing mutant neurons (threshold = 0.25) and (2) DLX-positive neurons identified by E2-crimson expression. Suite2p simultaneously exported ΔF/F₀ traces and spatial coordinates (x, y) for all identified neurons.

Exported datasets (ΔF/F₀ traces, neuronal coordinates, and DLX classification) were loaded into MATLAB for downstream analysis. ΔF/F₀ traces were deconvolved to infer single-action-potential-level neuronal activity, and spikes were detected using the “findpeaks” function. Spike frequency was calculated within specified time windows (expressed as spikes/minute). Network analysis was performed by treating deconvolved ΔF/F₀ signals as network nodes. Functional connections between neurons were quantified using Pearson correlation of deconvolved calcium traces, with connection strength defined by correlation coefficient magnitude and connection probability determined by thresholding correlated pairs Pairwise correlations between all neurons were compiled into correlation matrices. A threshold was applied to generate binary adjacency matrices, where edges represent significant functional connections between neurons^81^. Functional networks were constructed, where node positions were recalculated using force-directed layout algorithms (Kamada-Kawai algorithm^82^ and Fruchterman-Reingold algorithm^83^) based on pairwise correlation strength between neurons. This layout approach positions highly correlated neurons proximal to one another, thereby revealing functional clustering independent of spatial proximity. Network topological parameters were calculated for both representations and visualized accordingly.

### MosaicoidSim: Computational Framework

MosaicoidSim was developed as a computational framework to simulate organoid growth, cell migration, and spatial organization in mosaic contexts through sequential phases. The model was initialized with 1,000 cells in two-dimensional space within a circular domain. Each cell was assigned genotype identity (wildtype or MECP2-mutant), with initial cell positions randomly distributed or controlled according to experimental seeding conditions. We implemented a three-phase developmental model: (i) Initial cell seeding and aggregation along with embryonic body formation, (ii) proliferation and differentiation, and (iii) migration and maturation (**Fig. 3B**). The parameter of doubling time of cells at each stage was based on previous literature ^46–50^. The parameter of migration speeds was determined by previous experimental observations in cortical organoids ^33^. These phases respectively recapitulate: how initial cell positioning influences emergent spatial heterogeneity; how differential growth rates and differentiation efficiency between mutant and wildtype populations affect their relative proportions and spatial distributions; and how genotype-specific migration rates generate layer-specific neuronal organization characteristic of cortical development. The model started with 1,000 cells initialized in two-dimensional space, each assigned unique parameters reflecting empirically measured properties: genotype identity (wildtype or mutant), doubling time (proliferation rate), and cell adhesion factor (reflecting e.g. cadherin expression; **Fig. 3A**) ^46–50^. Cells were placed within a circular domain and sequentially repositioned according to their genotype-specific adhesion properties using Metropolis-type swap sorting (**Fig. 3B-i**). This stochastic approach recapitulates the homophilic adhesion preferences observed in real cells, where populations expressing similar cadherin types preferentially aggregate. To simplify without loss of generality, we assigned distinct cell-adhesion interaction parameters for homotypic (WT-WT, Mut-Mut) and heterotypic (WT-Mut) contacts, allowing the model to capture genotype-dependent clustering dynamics during the initial aggregation process that forms the embryoid body. During the proliferation and differentiation phase, cells grow heterogeneously, with outer cells proliferating faster and core cells more slowly due to oxygen diffusion limitations. To simulate this, we introduced a proliferation rate restriction parameter based on the local density gradient within organoids^84,85^ (**Fig. 3C**). Additionally, the decline in proliferation was modeled by incorporating a differentiation factor (τ) that accounts for oxygen diffusion limitations during growth. By tuning the τ value, which corresponds to the differentiation efficiency of cells, we could control the number of cells and organoid size by modulating the cell doubling time (**Fig. 3D, E**).

Phase 1: Initial Cell Seeding and Aggregation, Cells were positioned within the circular domain and sequentially repositioned according to genotype-specific adhesion parameters using Metropolis-type swap sorting. Distinct adhesion interaction coefficients were assigned for homotypic (WT-WT and Mut-Mut) and heterotypic (WT-Mut) cell contacts to simulate homophilic adhesion preferences and genotype-dependent clustering during embryoid body formation. Cells could reach equilibrium spatial distribution through iterative swaps minimizing adhesion energy. Phase 1 output: Final cell positions retained as input for Phase 2.

Phase 2: Starting from the equilibrium positions established in Phase 1, cells were assigned new parameters: genotype-specific doubling time (proliferation rate) and differentiation factor (τ). Heterogeneous proliferation was simulated by applying a proliferation rate restriction parameter based on local density gradients, with outer cells proliferating faster than core cells to recapitulate oxygen diffusion limitations. Differentiation efficiency was modeled by incorporating genotype-dependent τ values (τ_WT and τ_Mut) that controlled the rate of cell cycle exit and transition to non-proliferative states. Cell doubling time was modulated by the τ value to control final organoid size and cell population numbers. Phase 2 output: Updated cell positions and population composition retained as input for Phase 3.

Phase 3: Starting from the spatial distribution and cell composition established in Phase 2, we hypothesized WT:Mut migration rate difference of 10-fold based on MECP2 biology^33^. We systematically tested parameter combinations (WT: 0.5–2.0, Mut: 0.05–0.5 μm/h) and found WT=1.0, Mut=0.1 best reproduced experimental segregation patterns. cells were assigned new migration parameters: genotype-specific migration rates (e.g., WT: 1.0 μm/h; Mut: 0.1 μm/h). Radial migration was modeled such that each cell migrated outward from its current position toward peripheral regions according to its genotype-specific migration rate. Organoid space was classified into three concentric regions: core (innermost), intermediate (middle annulus), and peripheral (outer ring). Migration trajectories were calculated and tracked longitudinally across developmental timepoints (168, 288, 408, and 672 hours).

Systematic parameter sweeps were performed for Phase 1 (adhesion coefficients for WT-WT, Mut-Mut, and WT-Mut interactions), Phase 2 (differentiation efficiency values τ_WT and τ_Mut), and Phase 3 (migration rate parameters) to interrogate how genotype-dependent properties at each developmental stage collectively determine final spatial organization outcomes. *In silico* predictions were compared quantitatively to *in vitro* experimental organoids by comparing predicted cell distributions to fluorescently-labeled organoid imaging at matched timepoints. Simulations were implemented in Python and run for 100 independent iteration per condition.

### Quantitative analysis of degree of mosaicism

Spatial organization was quantified at each phase and timepoint using: (1) Segmentation Index (SI) ^86,87^, measuring genotype-based spatial clustering (SI < 0.5 indicating mixed distributions; SI > 0.8 indicating high segregation); (2) Moran’s Index ^52,53^, assessing spatial autocorrelation of genotype identity; (3) Mosaic Score, quantifying degree of mosaicism; (4) Radial Distance Analysis, calculating mean radial position of each genotype population from the organoid center; and (5) Regional Cell Stratification, quantifying the proportion of WT and Mut cells in core, intermediate, and peripheral regions.

### Reverse-transcription PCR

To assess gene expression profiles and biological activity in organoids, total RNA was extracted from organoid tissues using TRIzol reagent (Sigma) and further purified with the Direct-zol RNA Purification Kit (Zymo Research) according to the manufacturers’ protocols. Complementary DNA (cDNA) was synthesized from purified RNA using the SuperScript VILO cDNA Synthesis Kit (Thermo Fisher Scientific). Quantitative reverse transcription PCR (qRT-PCR) was performed on a QuantStudio 3 Real-Time PCR System (Applied Biosystems) using TB Green Premix Ex Taq II (Takara). Primer sequences for target and reference genes are provided in **Table S2**. GAPDH was used as the reference gene for normalization of relative mRNA expression. All qRT-PCR experiments were performed in triplicate using cDNA prepared from independent organoid pools (n ≥ 3).

### scRNA-seq and processing of transcriptional data

Eight organoids in each condition were dissociated with TrypLE express for 5-10 min. The cells were then washed with PBS + 0.1% BSA. Multiplexing oligos were incubated for 15 min and washed with PBS + 0.1% BSA three times. The cell suspension was then pooled into one tube and used to perform 10X Chromium followed by sequencing with NovaSeq. Raw sequencing reads were demultiplexed using demuxlet ^88^ to assign cell barcodes to individual samples and to identify and filter doublets. Demultiplexed FASTQ files were processed using the 10x Genomics CellRanger bioinformatics pipeline (v7.2) to generate count matrices. The CellRanger count function aligned reads to the human reference genome (GRCh38), performed cell barcode detection, and quantified gene expression across four experimental conditions: isogenic control organoids, MeCP2[R306C] mutant organoids, isogenic control organoids, and MeCP2[V247X] mutant organoids. EGFP reads were tracked as a transgene marker to identify and classify EGFP-positive (mutant) and EGFP-negative (control) cells within each condition. Count matrices from all four conditions were integrated as Seurat objects using Seurat (v5.0.1) and the Seurat wrapper integration function (v0.3.4) in R (v4.3.2), with EGFP expression status retained as a cell-level annotation for downstream analysis. Cells meeting the following quality control criteria were retained for downstream analysis: gene count ≥500 and ≤7000 genes per cell, and mitochondrial RNA content ≤20% of total transcripts. Low-quality cells and potential doublets not already filtered by demulet were removed based on these thresholds. Filtered count matrices from all four conditions were normalized using log transformation, scaled to unit variance, and integrated using the canonical correlation analysis (CCA) algorithm implemented in Seurat to correct for batch effects. Integrated data were centered per gene and underwent dimensionality reduction using principal component analysis (PCA). The cumulative variance explained by each principal component was visualized via elbow plot; the top 12 PCs (capturing ∼60% of variance) were retained for subsequent clustering analysis. Unsupervised clustering was performed using the shared nearest neighbor (SNN) algorithm with resolution parameter implemented in Seurat, followed by Louvain community detection with multilevel refinement to identify distinct cell clusters. Differentially expressed genes between clusters were identified using the Wilcoxon rank-sum test implemented in the FindAllMarkers function (log-fold-change threshold ≥0.25, adjusted p-value <0.05). Cell type annotation was performed by comparing cluster-specific gene signatures to known marker genes, including the following canonical markers: RGCs cells: *VIM, HOPX, ASCL1, OLIG2, EOMES, ASPM*. Excitatory neurons: *DCX, BCL11A, BCL11B, GRIA1, GRIA2, GRIN2B, GRIN2A, FOXG1, FOXP2, SATB1, SATB2*. Inhibitory lineage: *DLX1, DLX2, DLX5, DLX6, GAD1*, *GAD2*. Astrocytes: *S100B*, *GFAP*, and *APOE*. Based on these annotations, 5 different cell type populations were further clustered for sub-cluster analysis by recomputing the neighbor and dimensional reduction as well as UMAP plot for Pseudo-bulk RNAseq and pseudo-trajectory analysis. Ligand-receptor interaction analysis at the single cell level was performed using the CellChat R package (2.1.2) and NeuronChat ^89^ R package.

## Statistical analysis

The reported values are the means of a minimum of three independent experiments. Data are presented as the mean ± SD. For equal variances and normality distribution, a Student’s t-test was performed. To compare groups at multiple conditions, statistical comparisons were performed using one-way analysis of variance (ANOVA), with post-hoc pairwise comparisons carried out using the Tukey-Kramer method. Statistical tests were performed using GraphPad Prism 9 (GraphPad Software, San Diego, CA). p-values <0.05 or p-values <0.01 were considered significant in all cases.

## Data availability

All data supporting the conclusions are present in the paper and the supplementary materials. A reporting summary for this article is available as a Supplementary Information file. Read-level data from scRNA-seq (matrices, barcodes, and features), which have been deposited in the National Center for Biotechnology Information, Gene Expression Omnibus under accession code: GSE318601 and GSE318602. The processed data and imaging data generated in this study have been deposited in the Zenodo under accession code. Any additional information required to reanalyze the data reported in this paper is available from the corresponding author upon request.

## Code availability

The MATLAB code for quantifying neuronal activity data and R code for Mosaicoidseq, CellchatDB, and NeuroChatDB are available from GitHub. MosaicoidSim python code is available at Github repository (https://github.com/TatsuyaOsaki/MosaicoidSim). Source data are provided with this paper.

## Acknowledgments

We thank the Koch Institute’s Robert A. Swanson (1969) Biotechnology Center for technical support, specifically Microscopy, Flowcytometry core. We also thank Genome Technology Core at Whitehead Institute for Biomedical Research for Next-Generation sequencing on the Illumina for scRNAseq. We thank Evan Ruesch and Taylor Johns for editorial assistance, and members of the Sur lab for discussions and advice. This work was supported by NIH grants R01MH085802 and R01NS130361, MURI grant W911NF2110328, the Picower Institute Innovation Fund, and the Simons Foundation Autism Research Initiative through the Simons Center for the Social Brain to M.S.

## Author Contributions

T.O. conceived, designed, and performed the experiments. G.H. assisted with Femto3D Atlas anatomical scanning. T.O. analyzed the data, applied models to interpret the biological results, and generated the analyses and figures. T.O. and M.S. wrote the manuscript.

