## Supplemental figures and table for "Human cortical mosaicoids reveal impact on spatial organization and progressive loss of cross-genotype functional coupling in MECP2 mosaicism"

**Affiliations**

[†] Current affiliation

**This PDF file includes:**

Figures S1 to S10

Tables S1 to S2

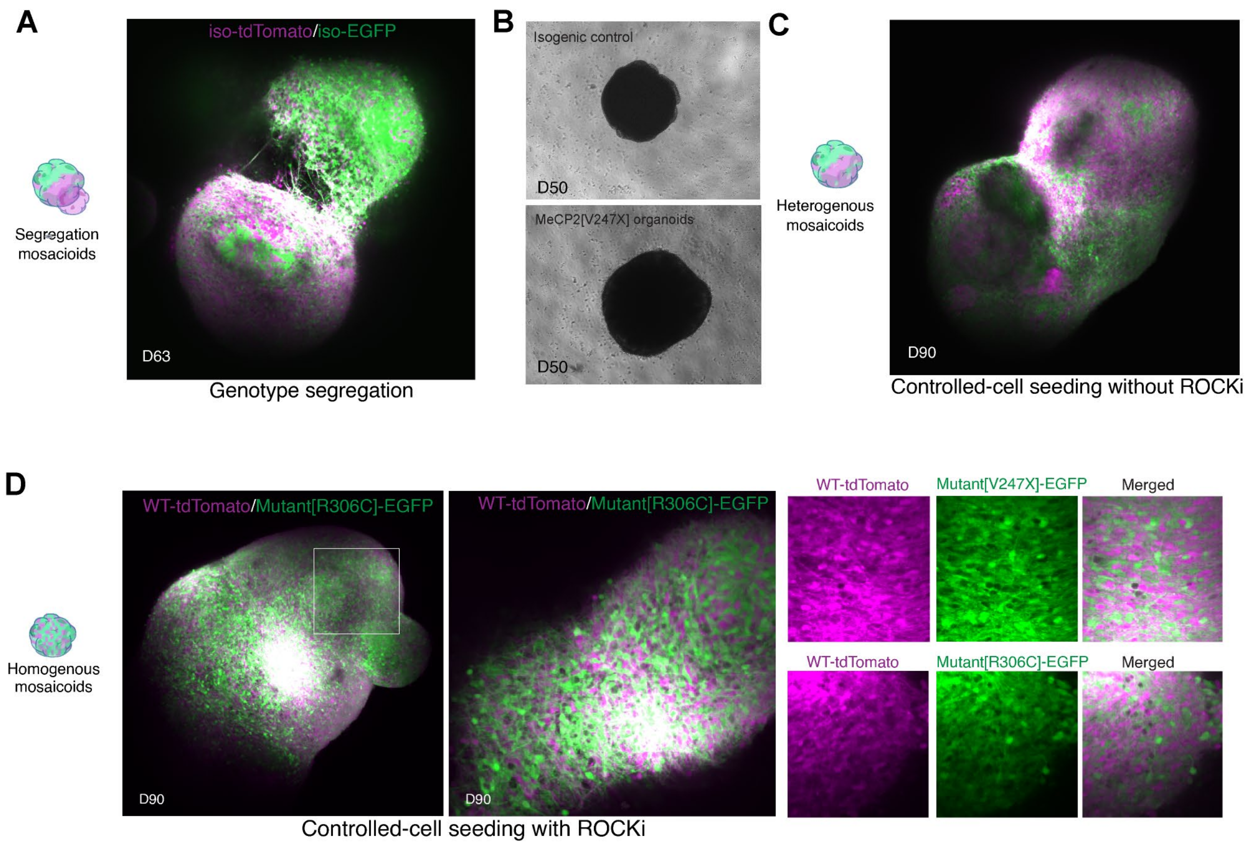


**Fig. S1| Controlled seeding and genotype identity determine the cellular architecture of cortical mosaicoids. (A)** Mosaicoid generated by fusing two organoids derived from genetically distinct donors [GM23279 and WIC07i-07982-4], imaged at day 63. The two populations occupy spatially distinct domains, indicating that donor-dependent differences in adhesive properties promote spatial separation. (**B**) Brightfield images of isogenic control (top) and MeCP2[V247X] mutant (bottom) cortical organoids at day 50 (D50), showing comparable morphology compared to mosaicoid formation. (**C**) Heterogeneous mosaicoid at day 90, generated by controlled single-cell seeding without ROCK inhibitor (ROCKi). WT-tdTomato and Mutant[R306C]-EGFP cells form irregular, variably sized clusters (**D**) Homogeneous mosaicoid at day 90, generated by controlled single-cell seeding with ROCKi. Left: low-magnification view of a WT-tdTomato / Mutant[R306C]-EGFP mosaicoid. Middle: magnified view of the boxed region. Right: high-magnification insets of WT-tdTomato with Mutant[V247X]-EGFP (top) and with Mutant[R306C]-EGFP (bottom), shown as single channels and merged, demonstrating uniform intermingling of the two genotypes in both mutation backgrounds.

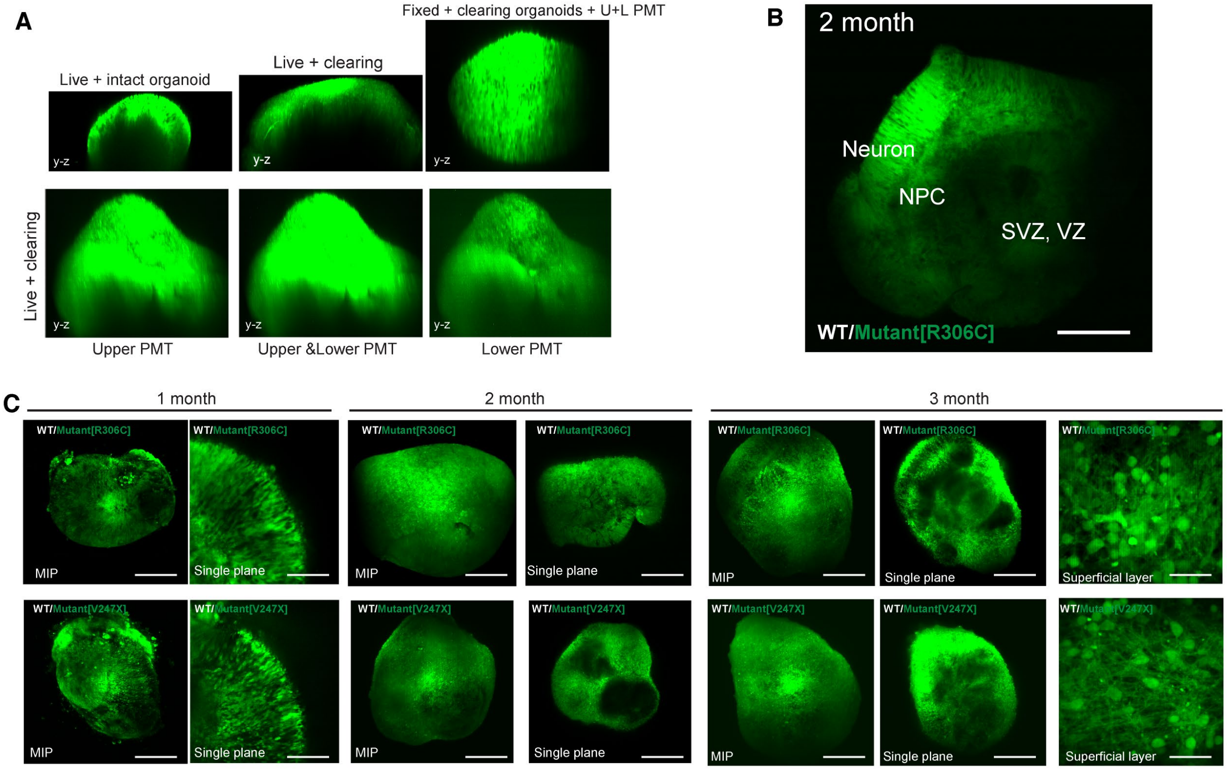


**Fig. S2| Dual-PMT collection system with refractive-index matching extends two-photon imaging to near-complete mosaicoid volumes and resolves genotype architecture across development. (A)** Y–Z cross-sections comparing imaging conditions and photomultiplier (PMT) configurations. Top row: live intact organoid (left), live organoid after iodixanol optical clearing (middle), and fixed, cleared organoid imaged with combined upper and lower PMTs (right). Bottom row: live cleared organoids imaged with the upper PMT alone (left), combined upper and lower PMTs (middle), and the lower PMT alone (right). The combined (U+L) configuration with optical clearing maximises photon collection and depth penetration. **(B)** Single-plane two-photon image of a 2-month WT / Mutant[R306C] mosaicoid. Cytoarchitectonic zones are identifiable from the surface inward: postmitotic neuronal layer, neural progenitor cell (NPC) zone, and subventricular/ventricular zone (SVZ/VZ). Scale bar, 500 µm **(C)** (C) Longitudinal volumetric imaging of WT / Mutant[R306C] (top) and WT / Mutant[V247X] (bottom) mosaicoids at 1, 2 and 3 months of differentiation. Maximum-intensity projections (MIP) and single planes are shown for each timepoint, with additional superficial-layer views at 3 months. Both views reveal progressive change in organoid size, internal architecture and mutant–wild-type spatial organisation over time, with V247X mosaicoids developing more pronounced genotype segregation. Scale bars, 500 µm.

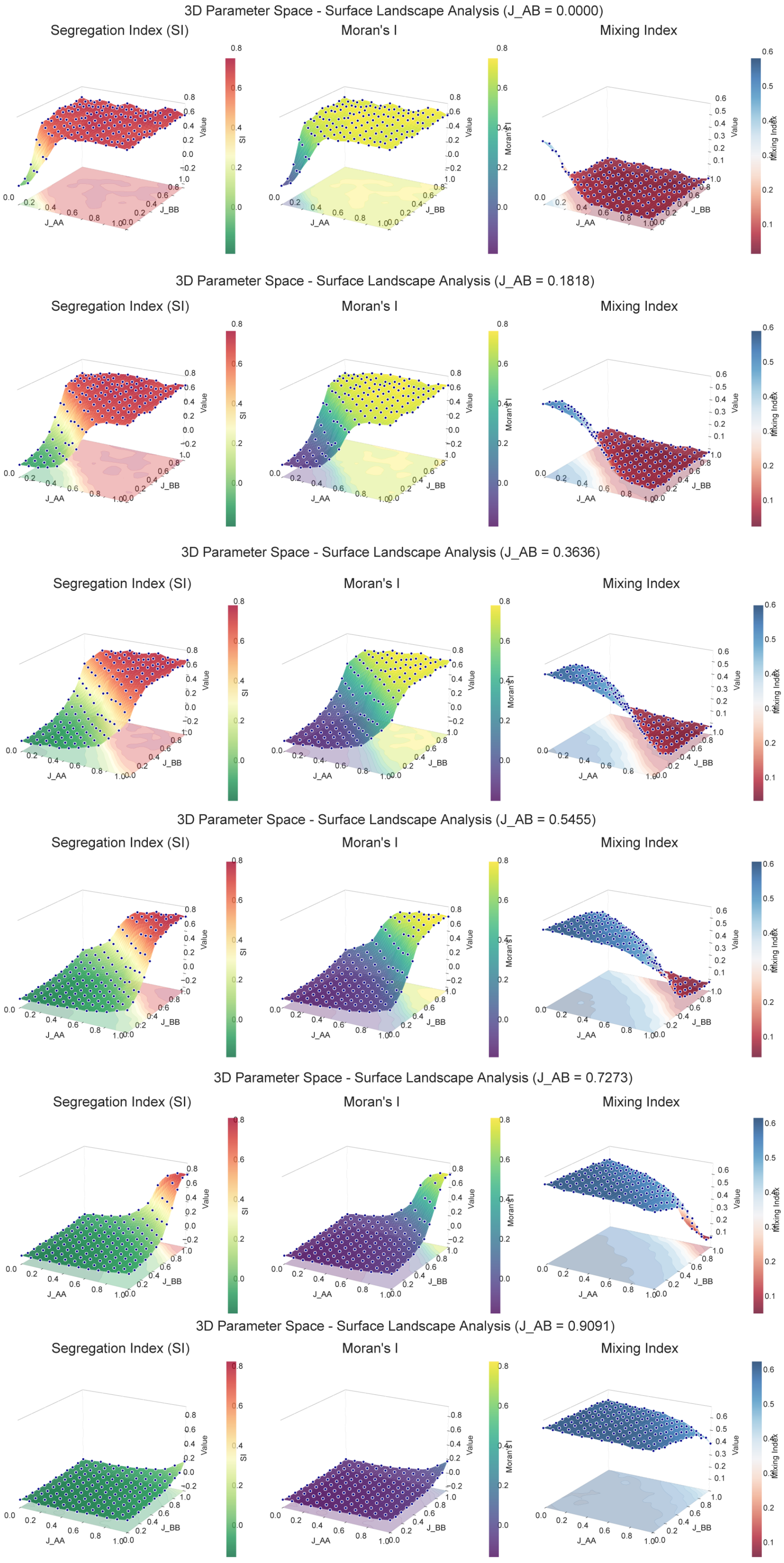


**Fig. S3| Heterotypic adhesion is the dominant determinant of spatial organisation across the full homotypic parameter space.**

In silico 3 dimensional surface landscapes of the Segregation Index (SI), Moran's I, and Mosaic Score as functions of the homotypic adhesion parameters J_AA (WT–WT) and J_BB (Mut–Mut), computed at six fixed values of the heterotypic parameter J_AB ([0, 0.18, 0.36, 0.55, 0.73, 0.91]; dimensionless interaction energies, see Methods). As J_AB increases, the SI and Moran's I surfaces collapse from high segregation toward uniform mixing across the entire J_AA × J_BB plane, while the Mosaic Score rises correspondingly. Heterotypic cell–cell interaction therefore sets the spatial outcome largely independently of homotypic adhesion. Each surface point is the mean of [100] independent simulations of 1,000 initial cells.

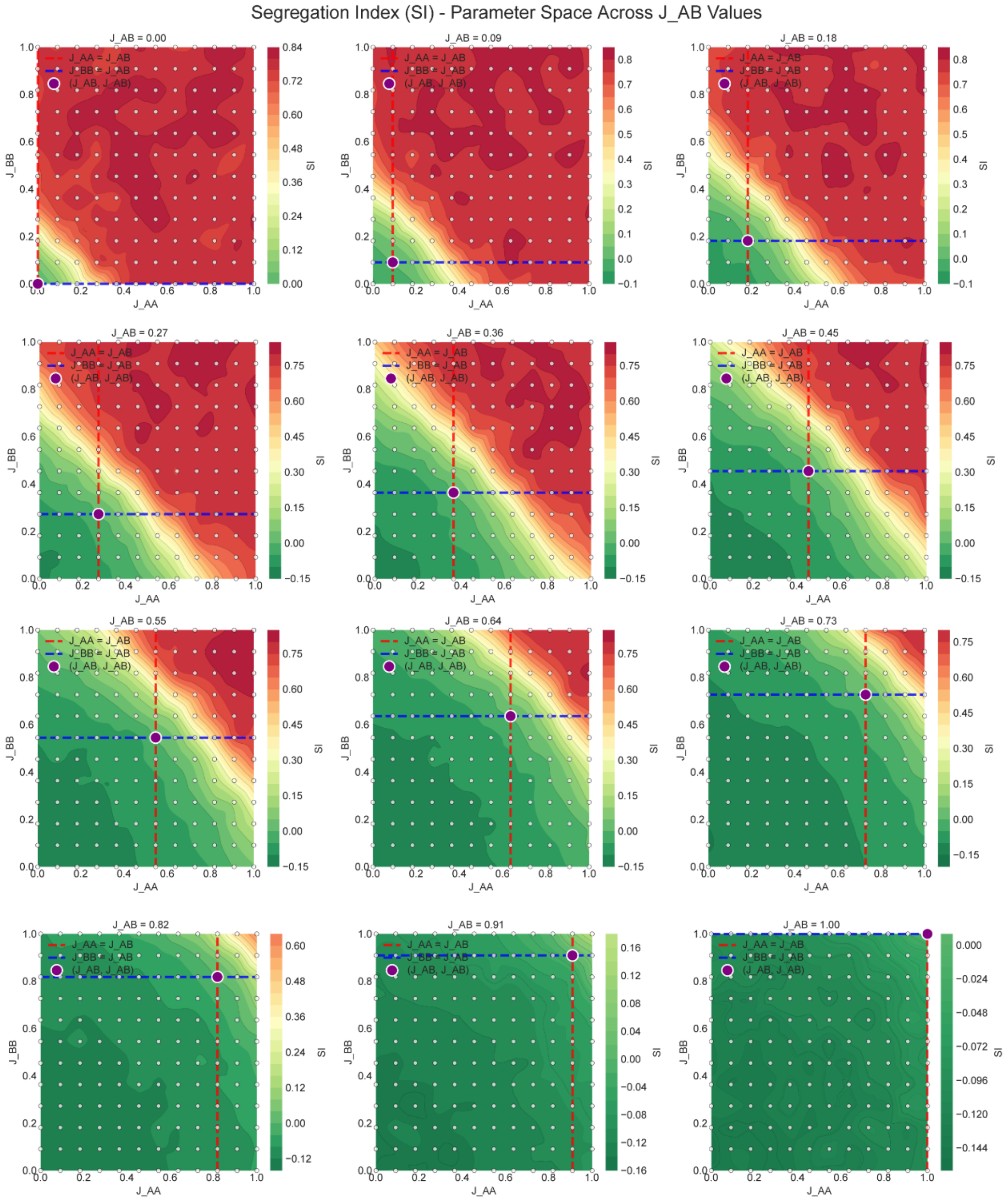


**Fig. S4| Segregation Index (SI) across the two-dimensional parameter space (J_WT-WT × J_MUT-MUT) maps confirm that segregation is lost above a threshold heterotypic adhesion.** Heatmaps of the Segregation Index (SI) over a grid of homotypic adhesion parameters J_AA (x-axis, WT–WT) and J_BB (y-axis, Mut–Mut), with panels arranged by increasing heterotypic adhesion J_AB from [0.00] to [1.00] (value given in each panel title). SI is defined as in Methods, where SI < 0.5 denotes a mixed distribution, SI ≈ 0.5 a random spatial distribution, and SI > 0.8 strong genotype segregation. Across the sweep, segregation persists over most of the homotypic plane at low J_AB but is abolished uniformly once J_AB exceeds the threshould, recapitulating the loss of segmentation observed on ROCKi treatment in vitro (Fig. 1E, Fig. S1D). Each cell is the mean of [100] independent simulations of 1,000 initial cells.(SI < 0.5 indicating mixed distributions; SI > 0.8 indicating high segregation)

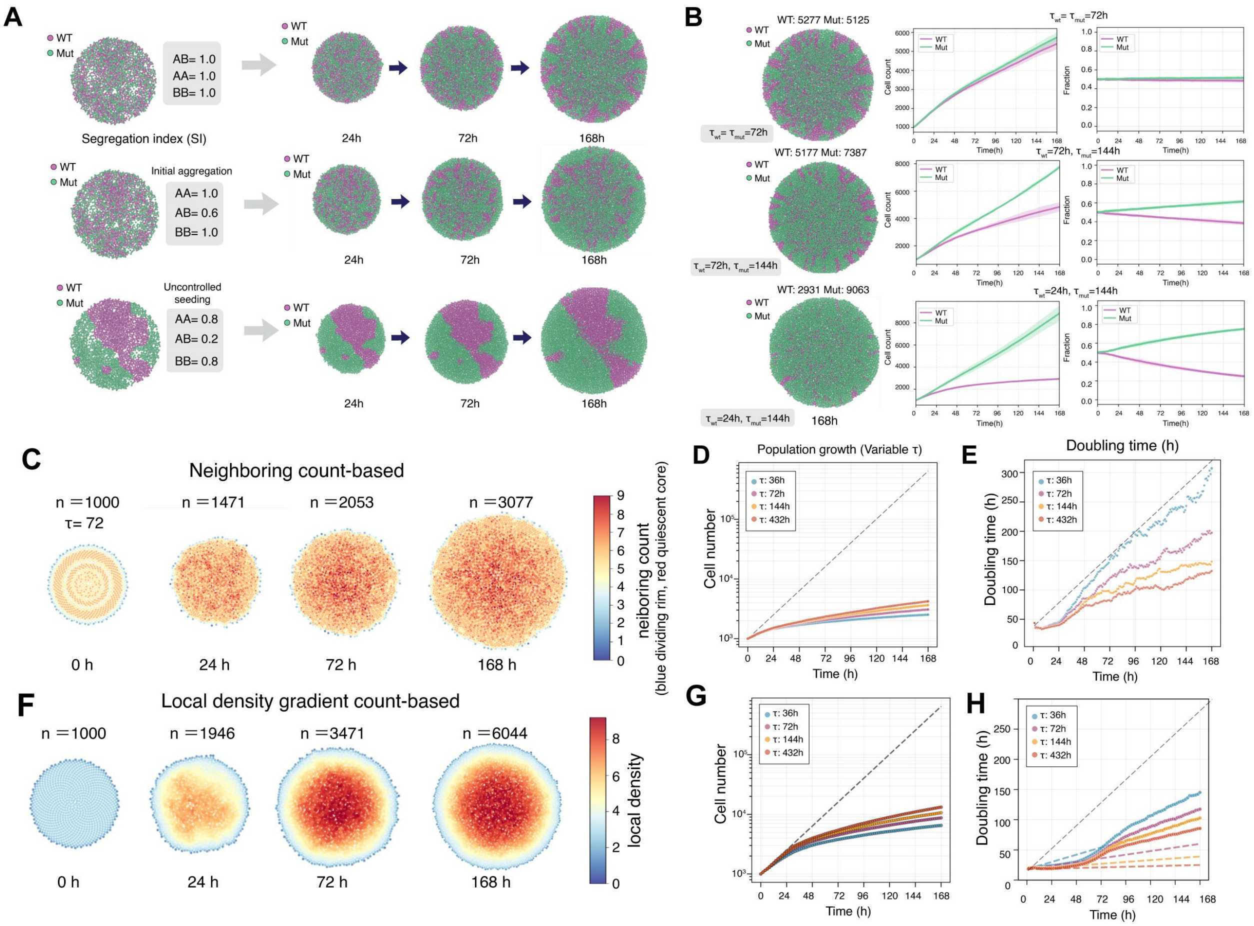


**Fig. S5| MosaicoidSim reproduces adhesion-driven phase separation, proliferation-driven clonal dominance, and density-dependent growth arrest. (A)** Simulated WT (magenta) and mutant (green) co-cultures at 24, 72 and 168 h under three adhesion regimes (J_AA, J_AB, J_BB as indicated), yielding mixed, partially segregated and fully phase-separated configurations. **(B)** Colony snapshots at 168 h (left), absolute cell counts (center), and fractional composition (right) for three combinations of WT and mutant doubling times (τ_WT, τ_Mut), illustrating how proliferative asymmetry drives clonal dominance. **(C)** Neighbor count heatmaps of a growing colony (τ = 72 h) at 0–168 h, revealing progressive expansion of a contact-inhibited quiescent core (red) relative to the proliferating rim (blue). **(D, E)** Population growth curves (D) and instantaneous doubling time (E) for intrinsic τ values of [36, 72, 144, 288, 432] h, showing density-dependent slowing of growth over time. **(F)** Local density-gradient maps of a simulated organoid at 0, 24, 72 and 168 h (*n* = 1,000 initial cells), illustrating the density field used to impose the proliferation-restriction parameter (Methods). **(G)** Total cell number over 168 h, showing exponential expansion before density-dependent arrest. **(H)** Simulated mean cell surface area over developmental time. All panels: 100 independent iterlation of simulations per condition; lines show the mean and shading the SD.

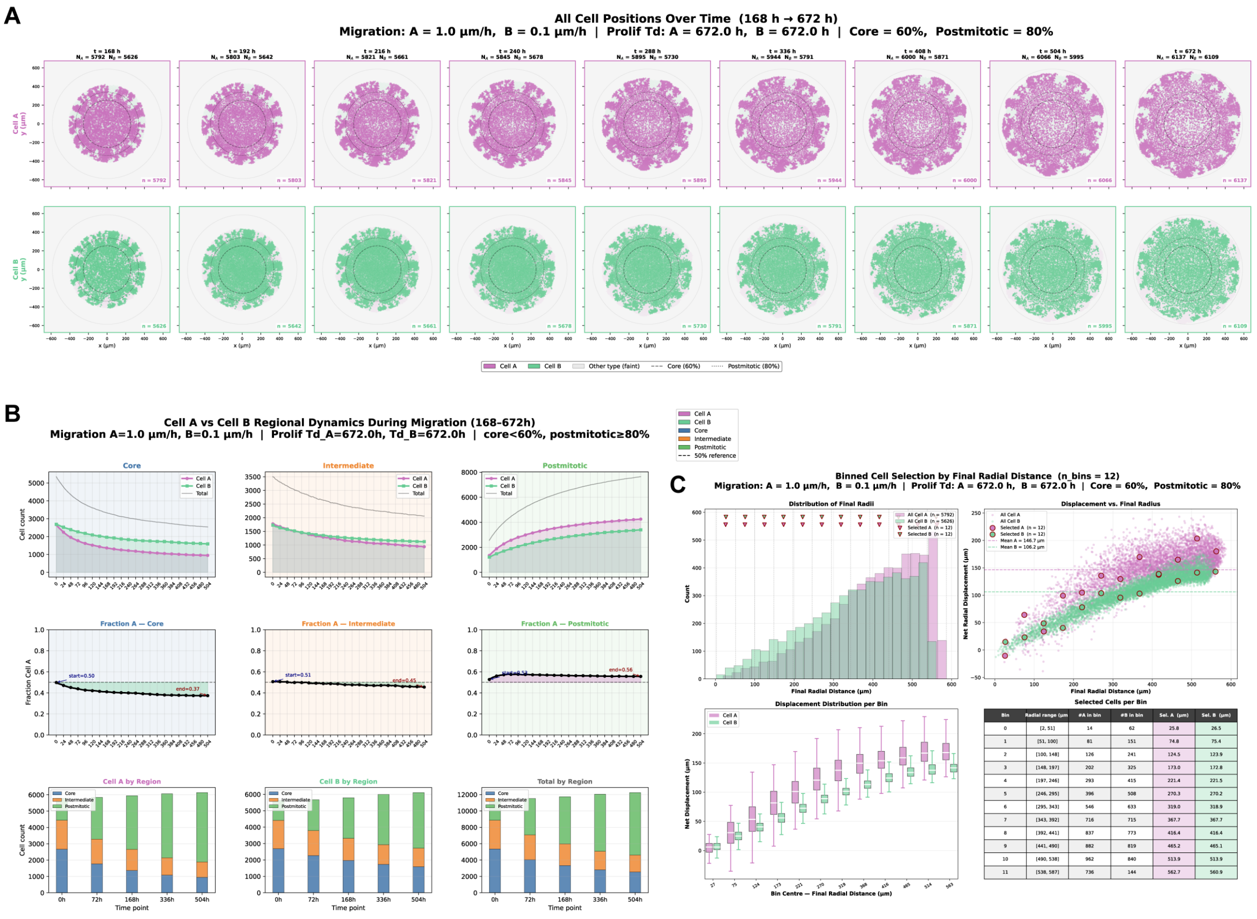


**Fig. S6| Differential migration rate alone generates radial genotype sorting in simulated R306C mosaicoids. (A)** Positions of WT (magenta) and mutant (green) cells from 168 to 672 h under asymmetric migration rates (WT 1.0 μm $h^{-1}$; Mut 0.1 μm $h^{-1}$) with matched doubling times (τ_WT = τ_Mut = 672 h), i.e. with proliferative asymmetry removed. Dashed circles mark the core (inner 60% of final radius) and postmitotic (outer 80%) boundaries. Faster-migrating WT cells disperse centrifugally while mutant cells remain confined to the core and intermediate zones. **(B)** Temporal dynamics of total cell count (top), WT fraction (middle), and regional composition (bottom) within the core, intermediate and postmitotic zones, for WT, mutant and total populations across five timepoints. **(C)** Binned analysis of final radial position (12 equal-width radial bins): cell-count distribution (top left), net radial displacement versus final radius (top right), per-bin displacement distributions (bottom left), and per-bin [mean ± SD of displacement and WT fraction] (bottom right). Radial sorting arises from migrational asymmetry alone. *n* = 1000 initial cells from 100 independent iternations.

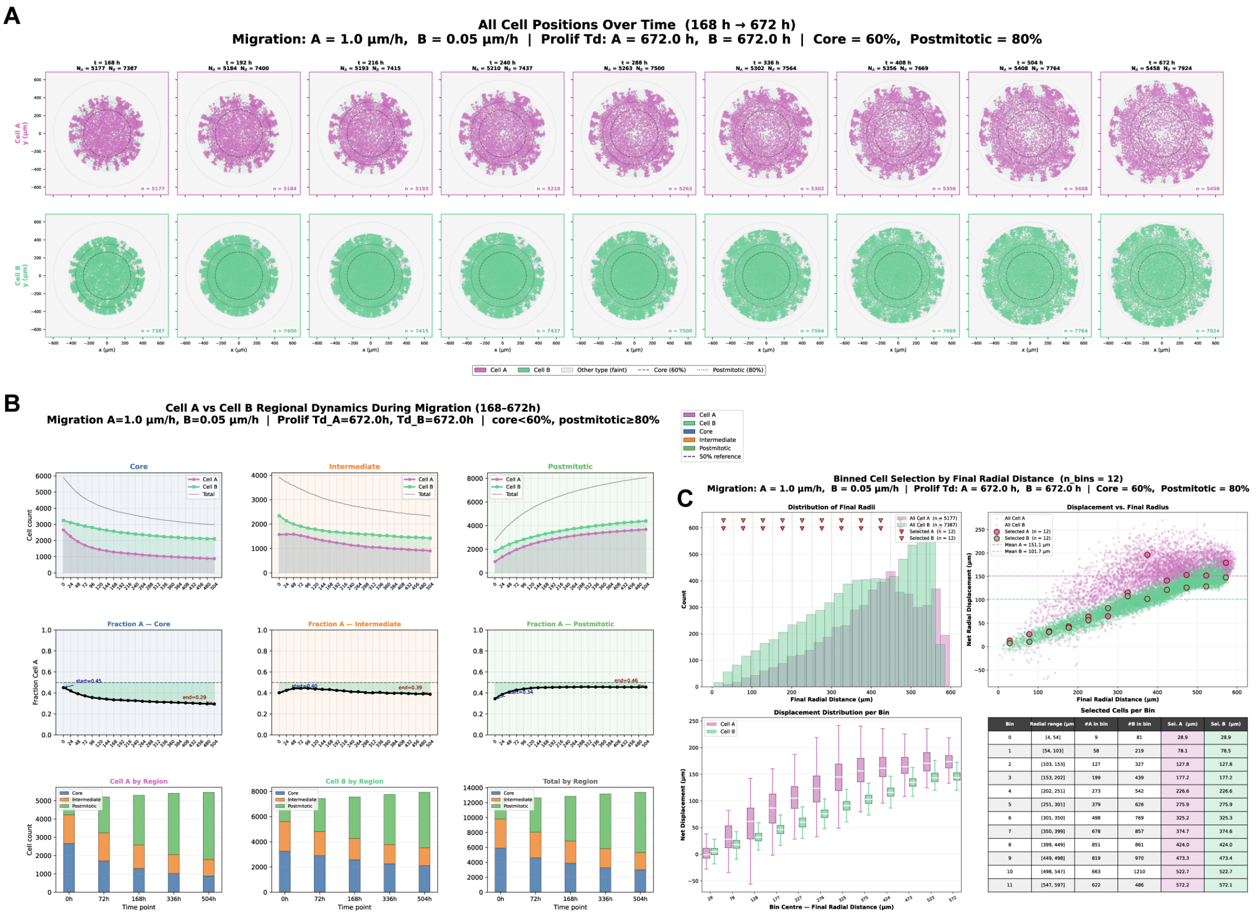


| Bin | Radial range (µm | #A in bin | #B in bin | Sel. A (µm) | Sel. B (µm) |
|---|---|---|---|---|---|
| 0 | [4, 54] | 9 | 81 | 28.9 | 28.9 |
| 1 | [54, 103] | 58 | 219 | 78.1 | 78.5 |
| 2 | [103, 153] | 127 | 327 | 127.8 | 127.8 |
| 3 | [153, 202] | 199 | 439 | 177.2 | 177.2 |
| 4 | [202, 251] | 273 | 542 | 226.6 | 226.6 |
| 5 | [251, 301] | 379 | 626 | 275.9 | 275.9 |
| 6 | [301, 350] | 498 | 769 | 325.2 | 325.3 |
| 7 | [350, 399] | 678 | 857 | 374.7 | 374.6 |
| 8 | [399, 449] | 851 | 861 | 424.0 | 424.0 |
| 9 | [449, 498] | 819 | 970 | 473.3 | 473.4 |
| 10 | [498, 547] | 663 | 1210 | 522.7 | 522.7 |
| 11 | [547, 597] | 622 | 486 | 572.2 | 572.1 |

**Fig. S7| Combined proliferative and migrational asymmetry produces severe superficial-layer mutant depletion in simulated V247X mosaicoids.** Parameters as in Fig. S6 except that the Phase 3 starting composition was set to the [70:30] mutant:WT ratio measured by flow cytometry in V247X mosaicoids (Fig. 1H) **(A)** Positions of WT (magenta) and mutant (green) cells from 168 to 672 h under asymmetric migration rates (A = 1.0 µm/h, B = 0.1 µm/h) with matched doubling times (672 h). Dashed circles denote the core (60%) and postmitotic (80%) zones. Faster-migrating Cell A progressively disperses outward while Cell B remains confined. **(B)** Temporal dynamics of cell counts (top), Cell A fraction (middle), and regional composition (bottom) within core, intermediate, and postmitotic zones for Cell A, Cell B, and total population across five time points. **(C)** Binned analysis (n = 12 bins) of final radial position showing cell count distributions (top left), net radial displacement versus final radius (top right), per-bin displacement distributions (bottom left), and summary statistics (bottom right), quantifying radial sorting driven by migrational asymmetry. $n$ = 1000 initial cells from 100 independent iternations.

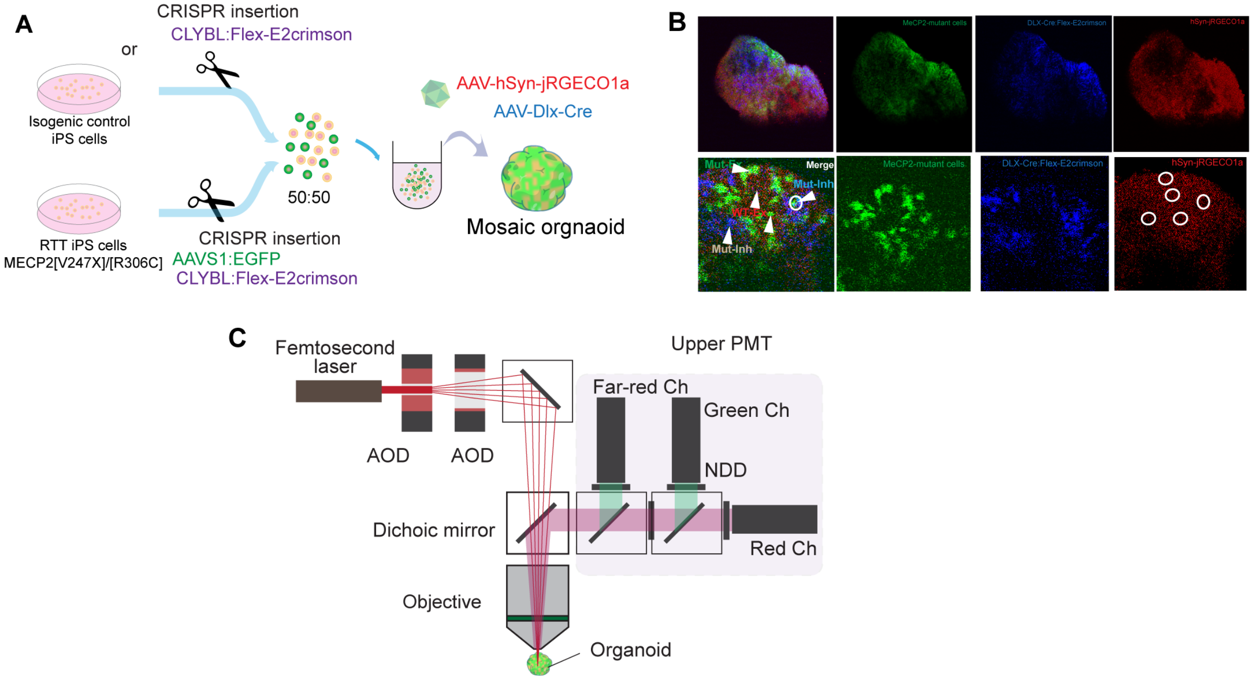


**Fig. S8| Experimental design for genotype- and cell-type-resolved calcium imaging of mosaicoids. (A)** Strategy for mosaicoid generation with genotype and phenotype readout. Isogenic control and RTT patient hiPS cells (MECP2[V247X] or MECP2[R306C]) carrying CRISPR-inserted reporters (AAVS1::CAG-EGFP in mutant cells; CLYBL::Flex-E2-Crimson) were FACS-deposited at a 50:50 ratio, then transduced with AAV-hSyn-jRGECO1a to report activity and AAV-hDLX-Cre to label inhibitory neurons; excitatory neurons were identified as jRGECO1a-positive, E2-Crimson-negative. (**B**) Representative images confirming cell-type-specific reporter expression: MECP2-mutant cells (EGFP, green), DLX-Cre-dependent Flex-E2-Crimson inhibitory neurons (far-red, displayed in blue), and hSyn-jRGECO1a (red). Scale bars, 50 µm. (C) Optical layout of the acousto-optic two-photon system: femtosecond laser (920 nm for EGFP, 1040 nm for jRGECO1a, 820 nm for E2-Crimson), four acousto-optic deflectors for random-access 3D scanning, dichroic mirrors, and three detection channels, green (540/50 nm), red (610/45 nm) and far-red (650 nm long-pass), read out through non-descanned detectors and the upper and lower PMTs for simultaneous multi-colour imaging.

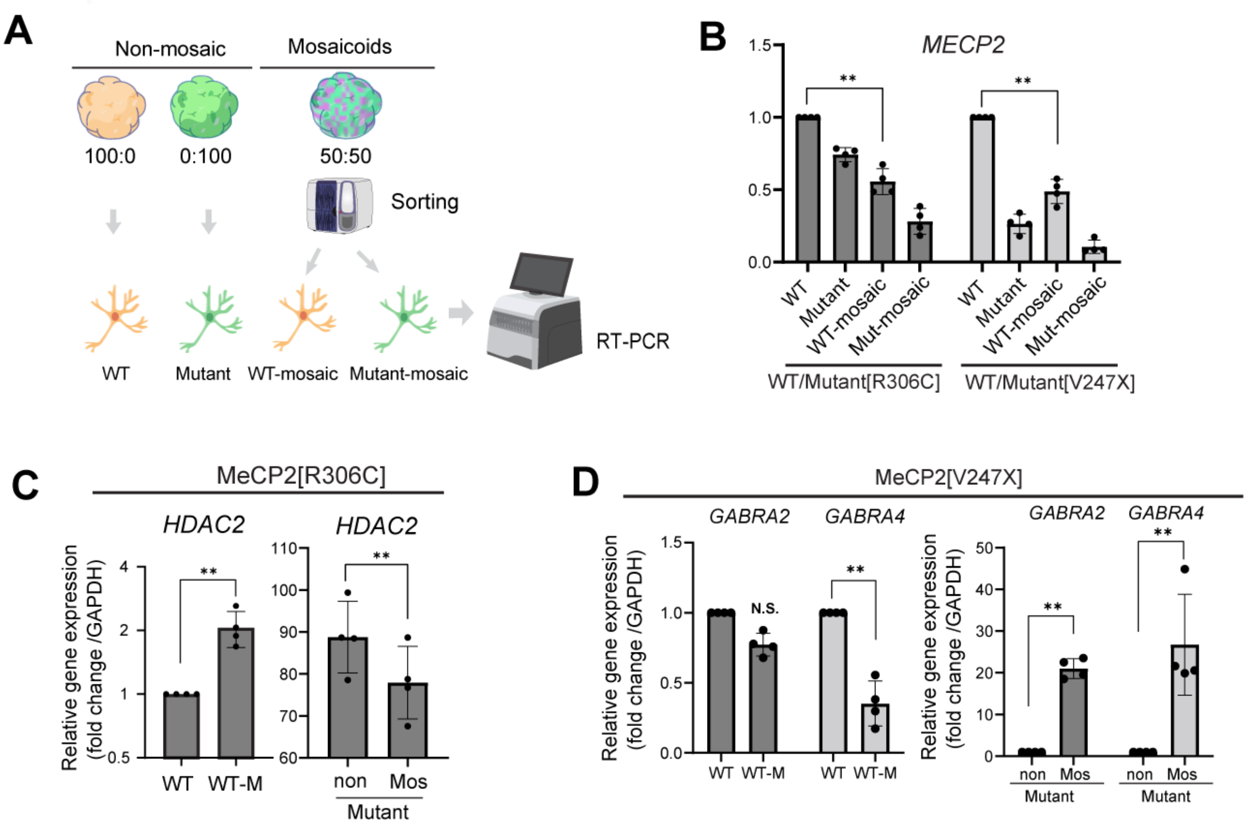


**Fig. S9| Cell-non-autonomous effect in mosaicoids.** (**A**) Mosaicoids and non-mosaic culture were generated as either 100% WT or 100% MECP2-mutant cells, alongside mosaicoids at a 50:50 ratio. WT- and mutant-derived cells were subsequently sorted from mosaicoids and analyzed by RT-PCR/qRT-PCR. (**B**) Relative MECP2 expression in WT, mutant, WT-mosaic, and mutant-mosaic cells carrying either the MECP2[R306C] or MECP2[V247X] mutation. MECP2 expression was reduced in mutant cells and further altered in the mosaic condition. (**C**) Relative HDAC2 expression in the MECP2[R306C] background. WT cells isolated from mosaicoids showed increased HDAC2 expression compared with non-mosaic WT cells, whereas mutant cells from mosaicoids showed reduced HDAC2 expression compared with non-mosaic mutant cells. (**D**) Relative expression of GABA receptor subunit genes in the MECP2[V247X] background. In WT-mosaic cells, GABRA2 expression was not significantly changed, while GABRA4 was reduced. In mutant cells, the mosaic condition significantly increased GABRA2 and GABRA4 expression compared with non-mosaic mutant cells. Expression levels were normalized to GAPDH. n = 4, mean ± SD; individual points indicate biological replicates (organoid). $^{**}p < 0.01$; N.S., not significant.

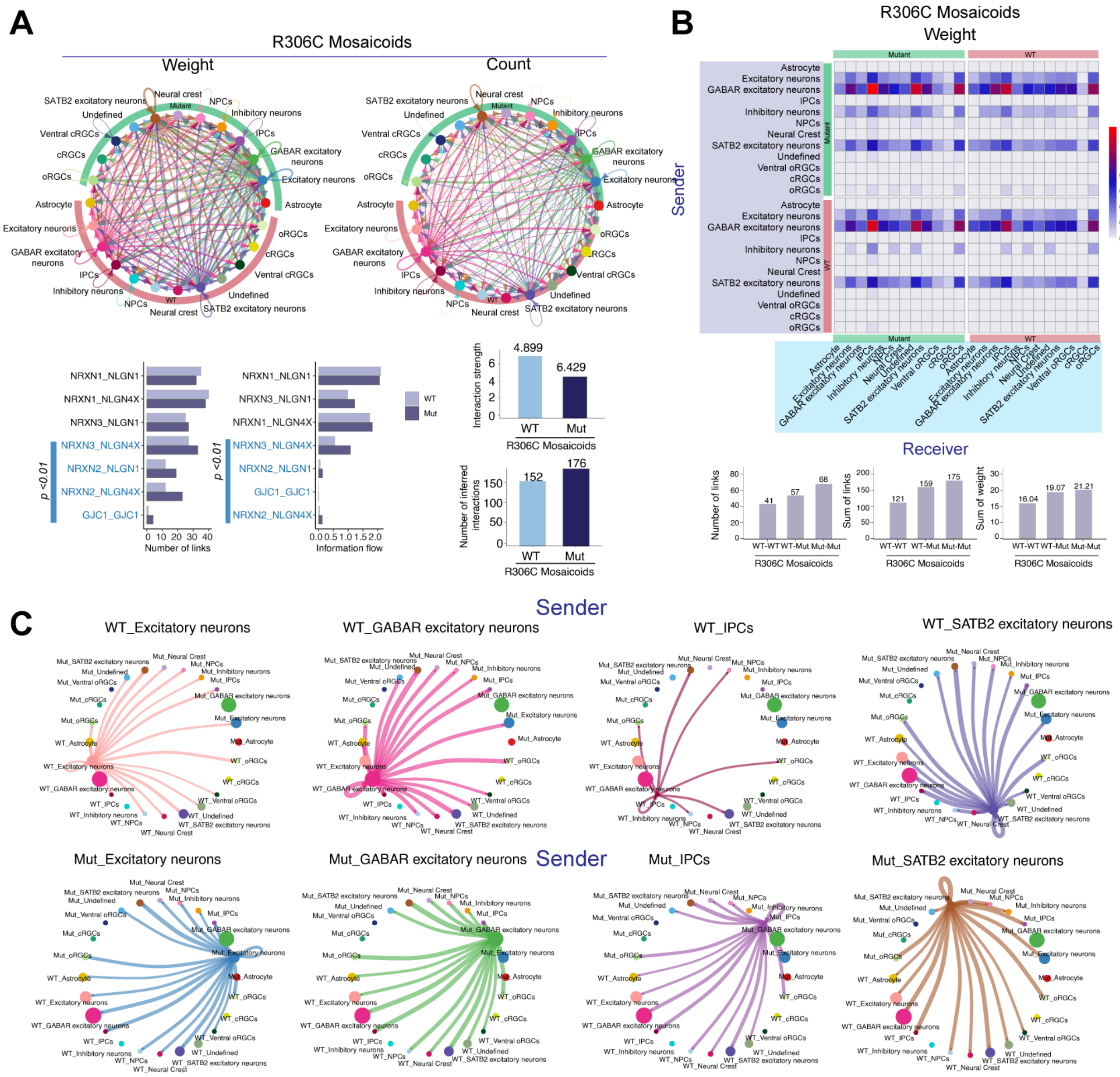


**Fig. S10| R306C mosaicoids retain an intact intercellular communication network with modest mutant–mutant over-connection. (A)** Global cell–cell communication networks inferred for R306C mosaicoids, summarized by total interaction weight and interaction count across annotated cell populations. Nodes represent cell types and edges represent predicted ligand–receptor-mediated interactions; edge thickness reflects interaction strength or number of interactions. Bar plots show the top altered signaling interactions ranked by number of links and information flow. Summary plots indicate increased overall interaction strength and number of inferred interactions in mutant mosaicoids compared with WT. **(B)** Sender–receiver heatmap showing predicted communication strength in R306C mosaicoids stratified. Rows indicate sender cell populations and columns indicate receiver populations, with color intensity representing normalized interaction weight. Bottom bar plots quantify the number and summed strength of interactions across WT and mutant mosaicoids. **(C)** Cell-type-specific outgoing signaling networks for selected sender populations in WT and mutant mosaicoids, including excitatory neurons, GABA-receptor-expressing excitatory neurons (GABAR excitatory neurons), intermediate progenitor cells, and SATB2 excitatory neurons. Curved edges indicate predicted outgoing interactions from each sender population to recipient cell types, highlighting genotype-dependent changes in communication patterns across neural and progenitor populations.

**Table S1. Key resources**

| Reagent and resource | Source | Identifier |
|---|---|---|
| **Cell line** | | |
| Rett syndrome patient derived iPS cells (female, 25Y, 705G deletion) | Coriell Institute | WIC07i-07982-4(MT) |
| Isogenic control pair, wild type | Coriell Institute | WIC07i-07982-2(WT) |
| Rett syndrome patient derived iPS cells (female, 8Y, missence) | Coriell Institute | WIC05i-127-325(MT) |
| Isogenic control pair, wild type | Coriell Institute | WIC04i-127-33(WT) |
| AAVpro 293T | Takara | N.A. |
| **Plasmid** | | |
| pAAV.CAG.GCaMP6f.WPRE.SV40 | Addgene | #100836 |
| pAAV.Syn.NES-jRGECO1a.WPRE.SV40 | Addgene | #100854 |
| pAAV-mDlx-GFP-Fishell-1 | Addgene | #83900 |
| pAdDeltaF6 | Addgene | #112867 |
| pAAV2/9n | Addgene | #112865 |
| pAAV2/1 | Addgene | #112862 |
| rAAV2-retro helper | Addgene | #81070 |
| pSpCas9(BB)-2A-GFP (PX458) | Addgene | #48138 |
| PX458-AAVS1 | Addgene | #113194 |
| AAVS1-tdTomato | Addgene | #159275 |
| AAVS1-Pur-CAG-EGFP | Addgene | #80945 |
| PX458-CLYBL | This study | This study |
| pC13N-iCAG.copGFP | Addgene | #66578 |
| CLYBL-Pur-Flex-E2Crimson | This study | This study |
| AAV-hDLX-Cre | This study | This study |

| **AAV** | | |
|---|---|---|
| AAV1. pAAV.CAG.GCaMP6f.WPRE.SV40 | Addgene | #100836-AAV1 |
| pAAV.Syn.NES-jRGECO1a.WPRE.SV40 | Addgene | #100854-AAVrg |
| pAAV-mDlx-GFP-Fishell-1 | Addgene | #83900-AAVrg |
| pAAV-hDLX-Cre | This study | |

**Table S2. qPCR primer**

| **Gene** | Forward primer 5'-3' | Reverse primer 5'-3' |
|---|---|---|
| **HDAC2** | ATG GCG TAC AGT CAA GGA GG | TGC GGA TTC TAT GAG GCT TCA |
| **MECP2** | GTG GAG TTG ATT GCG TAC TTC G | CCC TCT CCC AGT TAC CGT GAA |
| **GRABRA2** | TGC CCA ATG CAC TTG GAG G | GGA GCA ACC TGT ACT GAA TCA GA |
| **GRABRA4** | ACA ATG AGA CTC ACC ATA AGT GC | AAC TCC CGA ATT TCA AAG GGC |
| **GAPDH** | TGT GGG CAT CAA TGG ATT TGG | ACA CCA TGT ATT CCG GGT CAA T |
